# α-Synuclein acts as a cholesteryl-ester sensor on lipid droplets regulating organelle size and abundance

**DOI:** 10.1101/2024.06.19.599670

**Authors:** Reeba Susan Jacob, Alessandro Dema, Hélène Chérot, Calvin Dumesnil, Shira Cohen, Hadas Sar Shalom, Nitzan Rimon, Odelia Sibony-Nevo, Gilad Beck, Elena Ainbinder, Heimo Wolinski, Karin Athenstaedt, Francois-Xavier Theillet, Abdou Rachid Thiam, Philipp Selenko

## Abstract

While aggregated alpha-Synuclein (αSyn) is commonly associated with Parkinson’s disease, its physiological function as a membrane-binding protein is poorly understood. Here, we show that endogenous αSyn binds lipid droplets (LDs) in multiple human cell lines and in stem cell-derived dopaminergic neurons. LD-binding encompasses αSyn residues 1-100, which masks their detection by immunofluorescence microscopy, probably explaining the scarcity of similar observations in earlier studies. αSyn-LD interactions are highly temperature-sensitive and selective for cholesteryl-ester-rich LDs. They promote the formation of αSyn multimers that dissociate from LDs at non-permissive temperatures. αSyn remains LD-bound throughout starvation-induced lipolysis, whereas siRNA-knockdown diminishes LD abundance and compromises cell viability upon nutrient depletion, without affecting LD biosynthesis. Reciprocally, excess αSyn stimulates LD accumulation in dependence of lipid availability, restricts organelle size and ensures intracellular LD organization, which strictly depends on functional membrane-binding. Supporting a general role of αSyn in cellular lipid and cholesterol metabolism, our results point to additional loss-of-function similarities between Parkinson’s, Alzheimer’s and Gaucher’s disease.

## Introduction

Insoluble aggregates of human alpha-synuclein (αSyn) constitute the pathological hallmarks of Parkinson’s disease (PD) and related synucleinopathies, where they are found in remnants of midbrain dopaminergic neurons ^1^. For these reasons, efforts to unravel the cellular activities of αSyn mainly focused on possible mechanisms of aggregation ^2^ and how aggregated protein species may impose cytotoxic gain-of-function effects ^3^. By contrast, much less is known about the physiological role(s) of αSyn and whether loss-of-function scenarios contribute to disease development and/or progression. αSyn is abundantly expressed in different tissue and cell types, most prominently in midbrain neurons ^1^, but also in hematopoietic precursor cells ^4, 5^ and mature thrombocytes and erythrocytes ^6^. Currently, there is no common denominator to explain the presence of αSyn in these diverse sets of cells, nor about the biological activities it exerts in them. Because of their relevance for PD, possible functions of αSyn have been studied most extensively in nigrostriatal dopaminergic neurons, where high protein levels (∼40 μM) are found in presynaptic boutons ^7^. In these termini, αSyn colocalizes with synaptic vesicles (SVs) ^8^ and it has been implicated in SV binding and assembly ^9^, organization and trafficking ^10^, and general synaptic transmission ^11^. However, synuclein knockout mice reveal few impairments in these processes ^12, 13^ and no stably-associated αSyn was found on purified SVs ^14^, which highlights common shortcomings in confirming physiological membrane targets, contrary to the avidity with which αSyn interacts with reconstituted lipid vesicles *in vitro* ^15^.

Cellular αSyn primarily occurs as a cytoplasmic protein, with a smaller fraction in membrane-bound states that dynamically exchange with soluble αSyn ^16^. Different membrane compartments have been implicated in αSyn binding, including synaptic vesicles ^9^, lipid droplets ^17^, mitochondria ^18^ and the plasma membrane ^19^. In most instances, unique structural or compositional features were shown to mediate these interactions, including high curvature and corresponding lipid-packing defects in vesicles ^20^, polyunsaturated fatty acids and negatively charged phospholipids in planar membranes ^21^, or organelle-specific lipids such as cardiolipins in mitochondria ^18, 22^. These observations suggest that αSyn may interact with multiple different membrane compartments and probably based on their biophysical properties rather than their biological identities ^23^. Accordingly, metabolic activities, lipid availabilities and the inherent dynamics of membrane compositions and organizations are likely to modulate αSyn-membrane interactions, although there is no converging view about its function(s) on these membranes. Postulated activities range from the maintenance of membrane architecture to selective membrane-remodeling, including roles in membrane fission and fusion ^24^. Strikingly, cumulating evidence points to significant reciprocal effects that αSyn exerts on cellular lipid metabolism ^25, 26^, which strengthens the burgeoning notion that lipid-based defects play key roles in the pathologies of PD and related synucleinopathies ^16, 27, 28^.

Lipid droplets (LDs) are cellular organelles that store neutral lipids for general membrane homeostasis and energy production ^29^. Accordingly, LDs are found in cell types with active roles in lipid and cholesterol distribution, such as in glial cells of the central nervous system ^30^, for example, or in cells that primarily degrade lipids during mitochondrial *β*-oxidation, such as adipocytes in fat tissue ^31^. LD architectures are unique amongst cellular organelles in that they are surrounded by a single layer of membrane phospholipids, whereas their interiors are made up of neutral lipids in the forms of triacylglycerides (TAG) and cholesteryl-esters (CE) ^32^. TAG and CE contents of LDs vary considerably between cell types, often depending on whether they are used for energy production (i.e., TAG-rich, CE-poor), or membrane homeostasis (i.e., balanced TAG and CE) ^29^. LD proteins typically interact with the lipid monolayer in an integrated (Class I) or peripherally-associated fashion (Class II) ^33^, whereby the latter includes members of the perilipin protein family ^34^, which share sequence and structure similarities with synucleins ^35^. Ectopic expression of αSyn has been shown to trigger LD accumulation and binding in different cell types ^36–42^, although the nature of these effects and their biological relevance are unknown. Here, we demonstrate that αSyn acts as a cholesteryl-ester sensor on LDs and selectively interacts with CE-rich LDs to regulate their intracellular organization, abundance and turnover. By dissecting the molecular and biophysical determinants that govern the sterol-specificity of αSyn-LD interactions, we corroborate membrane-cholesterol homeostasis as a possible functional aspect of αSyn’s biology in health and disease ^43^.

## Results

To investigate interactions of αSyn with intracellular membranes, we electroporated recombinant, N-terminally acetylated protein into human A2780 cells ^44^. Within ∼10 h after αSyn delivery, we noted the formation of circular, cytoplasmic ‘exclusions’ (**Fig. 1A**) indicative of newly formed vesicles. Having ruled out that these structures corresponded to lysosomes (**Suppl. Fig. 1A**), we identified them as cellular lipid droplets (LDs) based on BODIPY staining and immunofluorescence colocalization with endogenous perilipin 2 (Plin2), a peripheral LD-marker protein ^45^ (**Fig. 1B, Suppl. Fig. 1B**). Surprisingly, we detected αSyn on LDs when we switched to antibodies against C-terminal αSyn epitopes, which revealed a prominent, rim-like decoration of LD-surfaces (**Fig. 1A, 1B**), reminiscent of published results of transiently overexpressed αSyn in non-related HeLa cells ^17^. By contrast, antibodies raised against N-terminal αSyn peptides up to residue 100 failed to detect the LD-bound protein and only displayed the homogenous distribution of predominantly cytoplasmic αSyn (**Fig. 1C, Suppl. Fig. 1C**). Thus, selective αSyn-detection on LDs coincided with the known structural boundary between helical, membrane-bound αSyn (aa 1-96) and its non-associated, disordered C-terminus (aa 97-140) on SDS micelles and reconstituted lipid vesicles ^46, 47^. These results suggested that αSyn interactions with artificial membranes and LDs involved a similar binding interface, whereby the latter rendered N-terminal epitopes inaccessible for antibody binding.

**Figure 1:**
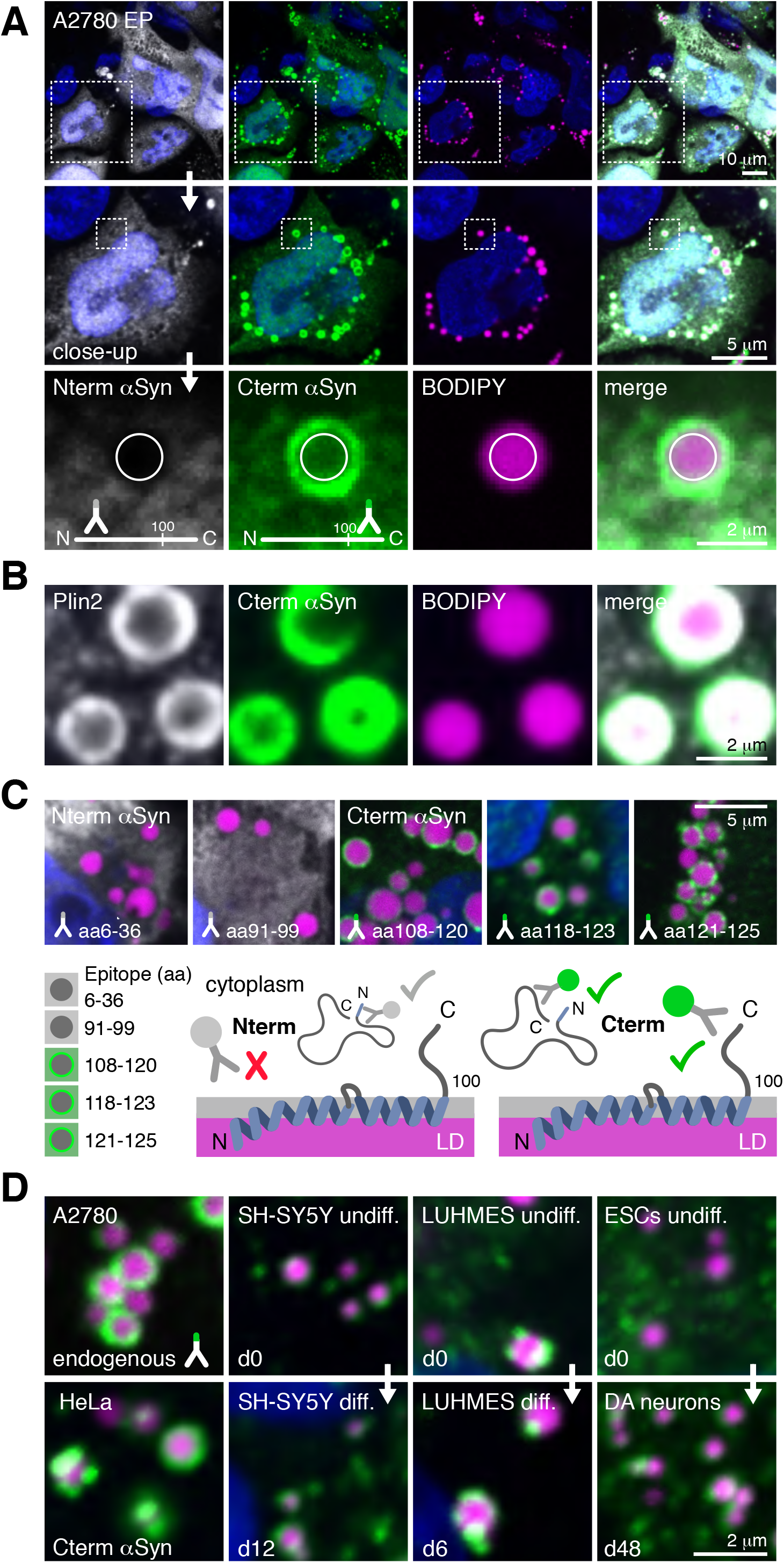
alpha-Synuclein (αSyn) binding to lipid droplets (LDs). (**A**) Electroporation (EP) delivery and immuno-fluorescence (IF) detection of intracellular αSyn with antibodies against N-(white) and C-terminal (green) epitopes in A2780 cells. Dashed boxes outline individual areas of magnification (top to bottom). BODIPY staining (purple) identifies cellular LDs. (**B**) IF-colocalization of endogenous perilipin 2 (Plin2, white) and EP-delivered αSyn (green) on LDs (BODIPY, purple). (**C**) Epitope-dependent IF-detection of EP-delivered αSyn on LDs. Amino-acid (aa) numbering indicates αSyn peptides used for monoclonal antibody production. Model representation of helical αSyn-LD interactions possibly giving rise to differential epitope accessibilities and selective αSyn detection. (**D**) LD-colocalization of endogenous αSyn in A2780 and HeLa cells, and in undifferentiated and differentiated SH-SY5Y and LUHMES cells, and in dopaminergic (DA) neurons derived from embryonic stem cells (ESCs) at the indicated days (d) of the respective differentiation protocols. For all IF images, cell nuclei/DNA are stained with DAPI (blue).

To rule out that differential antibody detection was an electroporation artifact, we transiently overexpressed αSyn in A2780 cells and found the same behavior (**Suppl. Fig. 1D**). Because naïve A2780 cells contain significant amounts of endogenous αSyn that localizes to different membrane compartments including the plasma membrane ^19^, we asked whether LD-colocalization was also observed in non-manipulated cells. Indeed, we detected αSyn on LDs of native A2780 and HeLa cells (**Fig. 1D, Suppl. Fig. 1E, 1F**), and in undifferentiated and fully differentiated neuronal SH-SY5Y and LUHMES cells (**Suppl. Fig. 1G, 1H, 1I**). Finally, we assessed αSyn expression and LD-association in human embryonic stem cells (ESCs) that we differentiated into midbrain-like dopaminergic (DA) neurons and co-stained with DA-specific markers such as microtubule-associated protein 2 (MAP2), class III beta-tubulin (Tuj1) and tyrosine hydroxylase (TH) ^48^ (**Suppl. Fig. 2A**). While LD abundance and αSyn expression were low at early differentiation time points, we observed colocalization of αSyn and LDs from day 31 onward (**Suppl. Fig. 2A**), including in TH-positive neurons at late stages of differentiation (day 48) (**Fig. 1D, Suppl. Fig. 2A**).

To confirm that αSyn interacted with existing and newly formed LDs in a similar manner, we first supplemented A2780 cells with increasing amounts of oleic acid (OA) to stimulate LD formation ^49^. Following, we delivered αSyn by electroporation and verified LD-colocalization (**Suppl. Fig. 2B**). In parallel, we electroporated αSyn before we treated cells with OA, aiming to generate new LDs in the presence of intracellular αSyn (**Suppl. Fig. 2C**). Both approaches revealed indistinguishable interactions between LDs and αSyn. Taking advantage of this effect, we purified LDs from OA-stimulated A2780 cells for *ex cellulo* binding assays with recombinant, AF488 fluorophore-labeled αSyn (N122C) (**Fig. 2A**). In line with what we found in intact cells, we detected avid interactions of wild-type αSyn with purified LDs (**Fig. 2B**). To determine whether LD-surface proteins contributed to αSyn binding, we treated purified LDs with trypsin, which efficiently degraded peripheral proteins such as Plin2, cytosolic phospholipase A2 (cPLA2), adipose triglyceride lipase (ATGL) and Rab5 ^45^ (**Suppl. Fig. 3A**). We detected no differences in binding to trypsin-treated or untreated LDs, which suggested that surface proteins did not contribute prominently to αSyn’s interaction. Focusing on direct membrane contacts instead, we first set out to delineate which parts of αSyn mediated LD-binding. Because our initial observations suggested that αSyn interactions with LDs and reconstituted lipid vesicles were similar, we probed possible contributions by membrane-anchoring residues ^50^ and found that N-terminally truncated (*Δ*N) and F4A-mutant αSyn failed to interact with purified LDs (**Fig. 2B**). We did not detect LD-binding of the PD-related, membrane-binding-compromised αSyn mutant A30P ^51^, whereas interactions of familial point-mutants A53T and E46K ^52^ were indistinguishable from the wild-type protein (**Fig. 2B**). Surprisingly, we found that Y39A αSyn did not interact with LDs (**Fig. 2B**), which identified this key aggregation-promoting ^53^ and postulated chaperone-homing residue ^54^ as essential for LD binding. We independently confirmed these results in intact A2780 cells (**Suppl. Fig. 3B**). In the course of these experiments, we noticed that αSyn did not interact with purified LDs when we put samples on ice before imaging (**Suppl. Fig. 3C**). Temperature-gradient experiments revealed that αSyn-LD binding was highly temperature sensitive and only occurred around 37 °C, but not at ambient temperatures of 25 °C and below (**Fig. 2C**, **Suppl. Fig. 3C**). By cycling the temperature between 25 °C and 37 °C, we learned that αSyn binding and unbinding was fully reversible (**Fig. 2D**). Furthermore, we found that increasing the temperature from 37 °C to 42 °C equally displaced αSyn from LD surfaces (**Fig. 2E**). These results suggested that temperature-sensitive LD-properties provided a contextual framework for αSyn interactions, with the known liquid-to-crystalline phase transitions of LD-core structures as likely candidates for the observed behavior ^55–57^.

**Figure 2:**
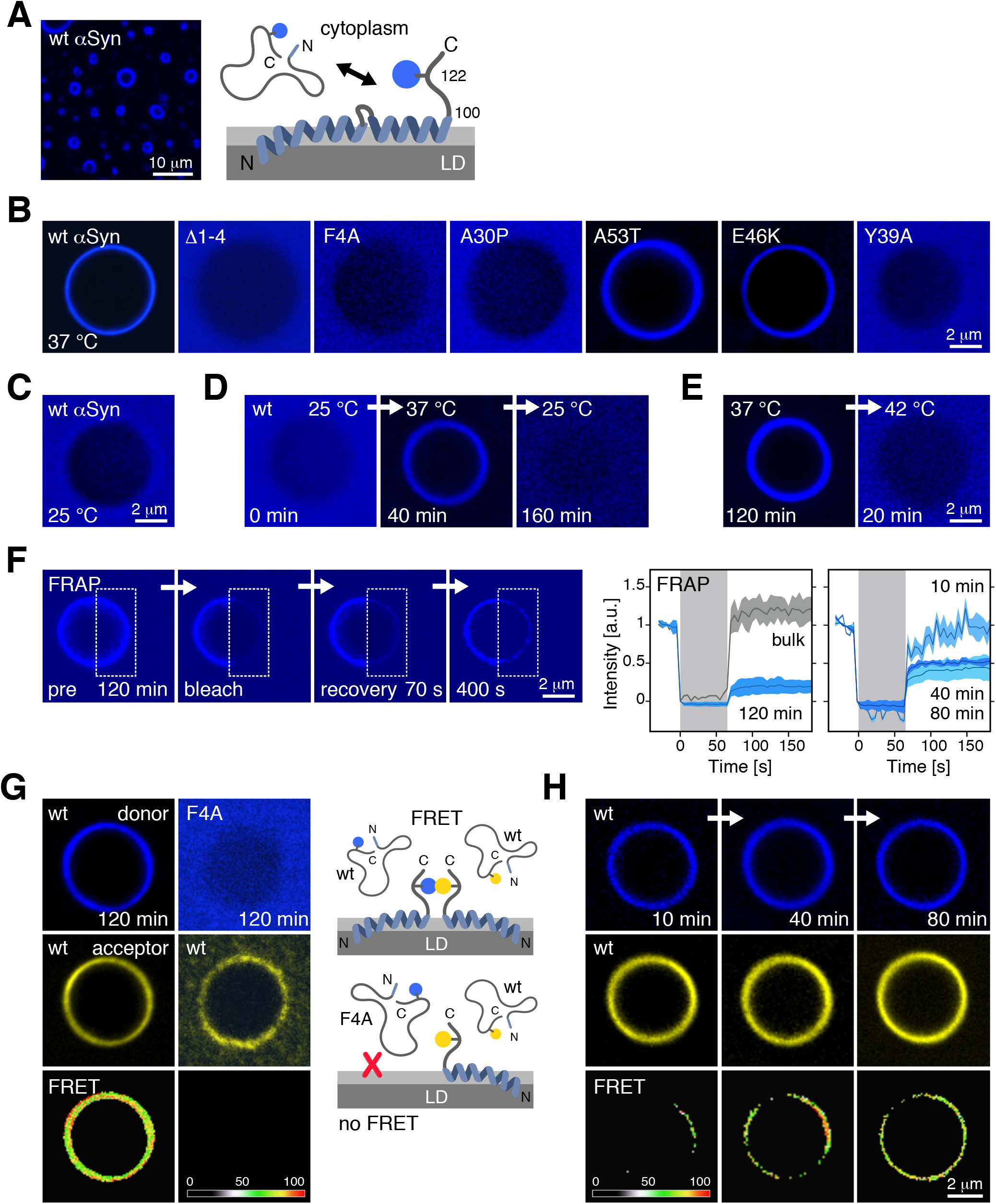
Biophysical characterization of αSyn-LD interactions. (**A**) Interaction of LDs purified from OA-supplemented A2780 cells with Alexa Fluor 488-labeled (AF488, pseudo-colored in blue), recombinant αSyn. The helical αSyn model depicts the C-terminal fluorophore position in the engineered asparagine to cysteine (N122C) mutant. (**B**) Binding of AF488-labeled wild-type (wt) and indicated mutant forms of αSyn to purified LDs at 37 °C. All mutations were engineered into the αSyn N122C variant. (**C**) Loss of LD-binding of AF488-labeled wt αSyn at 25 °C. (**D**) Selected time-points of temperature-cycle experiments between 25 °C and 37 °C, and (**E**) between 37 °C and 42 °C indicate reversible αSyn-LD association and dissociation within a narrow temperature range. (**F**) Exemplary fluorescence images of wt AF488-labeled αSyn on LDs pre and post photo-bleaching (120 min of incubation at 37 °C). Photo-bleached areas are outlined in dashed lines. Fluorescence-recovery after photobleaching (FRAP) traces (left panel, blue) and comparison with unbound αSyn in buffer (bulk, dark grey). Extended photobleaching (∼70 s) was used to efficiently ablate AF488 fluorescence (shaded grey area). Corresponding FRAP traces of wt αSyn after LD-incubation for 10 (light blue), 40 (dark blue) and 80 min (cyan) at 37 °C (right panel). (**G**) Förster energy-transfer (FRET) experiments of donor (AF488) and acceptor (AF596) fluorophore-labeled wt αSyn incubated for 120 min with purified LDs at 37 °C (left panel) and of a wt/F4A-mutant αSyn mixture under the same experimental conditions (right panel). Top panels depict donor and acceptor fluorescence intensities on LDs. FRET efficiencies are color-coded along a scale of lowest and highest signals (bottom panels). (**H**) Time-evolution of wt αSyn-αSyn FRET on individual LDs at 37 °C. Uniform donor and acceptor fluorescence are detected before FRET-positive species form.

To study the dynamics of αSyn-LD association and dissociation in more detail, we performed fluorescence-recovery after photobleaching (FRAP) experiments on purified LDs that we incubated with AF488-labeled αSyn. At 37 °C and incubation for 120 min, we only recovered a fraction of the initial fluorescence after photobleaching (**Fig. 2F**). Notably, we obtained increasingly higher fluorescence recovery yields when we shortened LD-incubation times (**Fig. 2F**), whereas the initial slopes of individual recovery rates were similar (**Suppl. Fig. 3D**). From these results, we hypothesized that LD-bound αSyn matured into more stably associated and less diffusive species, unable to exchange with bulk αSyn (**Suppl. Fig. 3E**). Accordingly, the difference in recovered fluorescence corresponded to the fraction of αSyn molecules in these slowly dissociating macrostructures (**Fig. 2F, Suppl. Fig. 3E**). Because αSyn has been shown to oligomerize on LDs ^17^, we reasoned that αSyn multimers or aggregates represented obvious candidates for such dark-state species ^58–60^. To determine whether reduced LD-dissociation correlated with αSyn oligomerization, we performed Förster resonance-energy transfer (FRET) experiments with mixtures of donor (AF488) and acceptor (AF594) fluorophore-labeled αSyn that we added to purified LDs. At 37 °C and incubation for 120 min, we observed strong FRET intensities, which confirmed the formation of αSyn multimers (**Fig. 2G**). To rule out that FRET signals originated from αSyn molecules that spontaneously aggregated on or off LDs, we repeated these experiments with a mixture of wild-type and membrane binding-deficient F4A-mutant αSyn (**Fig. 2B**). We did not detect FRET signals under these conditions (**Fig. 2G**), which confirmed that αSyn oligomerization was contingent on functional membrane-binding. Next, we asked whether FRET signal-intensities increased with LD-incubation time, as predicted by our model. While we observed uniform donor and acceptor fluorescence on LD surfaces already after 10 min of incubation, we only detected few instances of FRET, whereas signals increased and became more continuous upon prolonged incubation (**Fig. 2H**), reminiscent of the seeding-induced growth behavior of canonical αSyn aggregates ^61, 62^. To investigate early steps of LD-mediated αSyn-αSyn interactions, we followed the evolution of FRET signals in wide-field mode, which allowed us to monitor multiple LDs simultaneously. Surprisingly, we found that FRET-active species randomly formed on individual LDs and often in multiple, discrete spots independent of LD size and curvature (**Suppl. Fig. 3F**). In many instances, we detected FRET signals on large LDs but not on neighboring specimens of smaller dimensions (**Suppl. Fig. 3F**), which suggested that αSyn oligomerization on LDs likely followed a stochastic nucleation process. These FRAP and FRET results demonstrated that αSyn sampled different structural and dynamic states upon LD-binding, which collectively converged into more stably-associated αSyn multimers.

To further unravel the molecular basis of αSyn-LD interactions, we set out to dissect possible contributions by different LD components ^32^. Because LDs lack many of the core ingredients of prototypic αSyn-membrane interactions, such as negatively-charged phospholipids ^63^ or appreciable curvature effects given their large size ^20^, we first focused on membrane surface constituents and asked whether LD-specific lysophospholipids (LPLs) and LPLs-dependent packing defects contributed to αSyn binding, as proposed recently ^64^. To this end, we treated A2780 cells with bromoenol lactone (BEL), methyl-arachidonylfluorophosphonate (MAFP), or pyrrolidine-2 (Py-2) to inhibit biosynthesis of the major LPL lysophosphatidylcholine (see **Materials & Methods** for details) (**Suppl. Fig. 4A-D**). Neither of these compounds affected colocalization of electroporation-delivered αSyn with LDs. Next, we diminished LD-TAG contents by inhibiting non-specific long chain acyl-CoA synthetase (ACSL) with Triacsin C (TriC) and diacylglycerol transferase 1 (DGAT1) and 2 (DGAT2) with T863 and PF-06424439, respectively (**Suppl. Fig. 4E-G**). αSyn-LD interactions were unaffected. Finally, we probed CE contributions to αSyn LD-binding by inhibiting lysosome function with bafilomycin A1, which blocks lysosomal acidification and, in turn, the ability to distribute cholesterol for CE synthesis. Having confirmed the intended drug activity by loss of pH-sensitive lysotracker staining in treated A2780 cells, we found that αSyn no longer bound to LDs (**Fig. 3A, Suppl. Fig. 5A**). Similarly, we detected no binding of αSyn to cellular LDs when we inhibited the lysosomal cholesterol transporter NPC1 with U18666A (**Fig. 3A, Suppl. Fig. 5B**). Providing a third line of evidence in support of CE as the key determinant for αSyn-LD interactions, we showed that direct inhibition of the CE-synthesizing acylCoA:cholesterol acyltransferase (ACAT) with Avasimibe equally abolished αSyn-binding to LDs (**Fig. 3A, Suppl. Fig. 5C**). To further substantiate these conclusions, we aimed at rescuing αSyn-LD interactions under persistent bafilomycin exposure by direct addition of cholesterol to the growth medium of lysosome-inhibited A2780 cells. Indeed, bypassing lysosomal function via the external supplementation with cholesterol restored αSyn-LD interactions (**Fig. 3A, Suppl. Fig. 5D**). Together, these data provided strong evidence that the CE component of LDs acted as the determining factor for αSyn binding.

**Figure 3:**
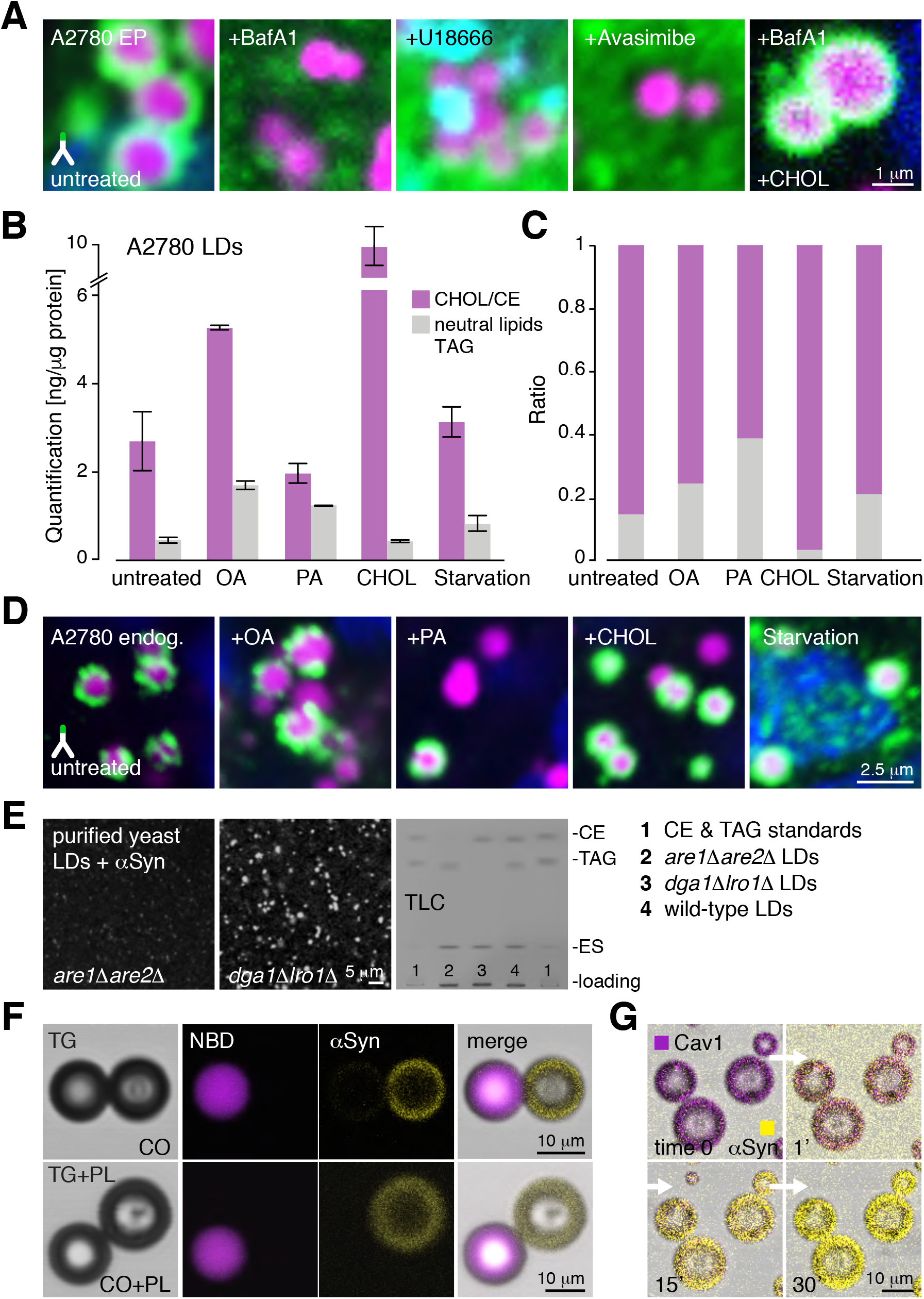
Sterol levels determine αSyn-LD binding. (**A**) IF-microscopy closeup views of EP-delivered wt αSyn (green) in A2780 cells and colocalization with LDs (purple) upon treatment with Bafilomycin A1 (BafA1), U18666, Avasimibe and BafA1 with direct cholesterol (CHOL) supplementation. (**B**) Colorimetric quantification of triacylglyceride (TAG) and cholesteryl-ester (CE) contents of LDs purified from A2780 cells grown in complete medium (CM), or in the presence of oleic acid (OA), palmitic acid (PA), cholesterol (CHOL) and under starvation. Bar graphs denote TAG and CE levels normalized to total protein concentrations, represented as mean ± SD from two independent batches of purified LDs. (**C**) Corresponding CE:TAG ratios of purified LDs. (**D**) IF-microscopy closeups of endogenous αSyn and colocalization with LDs in A2780 cells grown in the presence of OA, PA, CHOL and under starvation. For all IF images, cell nuclei/DNA are stained with DAPI (blue). (**E**) Fluorescence microscopy detection of AF488-labeled aSyn incubated at 37 °C with LDs purified from yeast strains devoid of TAG (*lro1*Δ*dga1*Δ) or sterol esters (*are1*Δ*are2*Δ), respectively. Thin layer chromatography (TLC) analysis of whole-cell lipid extracts from wild-type and mutant yeast strains, with ergosterol, TAG and CE in the lipid reference mixture. (**F**) Mixtures of pure droplet emulsions made up of cholesterol oleate (CO) and NBD-labeled Triolein (purple, top-row) or of CO and NBD-Triolein emulsions surrounded by phospholipids (PL, bottom row) incubated with AF594-labeled αSyn (AF594, pseudo-colored in yellow) at 37 °C. (**G**) Cholesteryl-oleate droplets pre-incubated with the amphipathic helix (AH) of caveolin 1 (Cav1)-labeled with NBD (pseudo-colored in purple). Time-resolved displacement of AH-Cav1 upon addition of AF594-αSyn to the bulk medium at 37 °C.

To further corroborate CE’s role in mediating αSyn association, we quantified TAG and CE levels in LDs that we purified from A2780 cells grown under different media conditions. In complete medium (CM), A2780 LDs contained a roughly threefold excess of CE over TAG in nanograms (**Fig. 3B**) leading to a CE:TAG ratio that was significantly skewed towards the sterol component (**Fig. 3C**). Given the importance of CE in αSyn binding, this cell type-specific imbalance agreed well with the observed avidity with which αSyn interacted with A2780 LDs. Upon supplementing growth media with OA, we measured a uniform increase of TAG and CE levels in LDs, which did not significantly alter the overall ratio between the two components (**Fig. 3B, C**), in line with the equally good binding behavior of αSyn to LDs obtained from OA-supplemented cells (**Fig. 1D**, **Fig. 2**). By contrast, we determined significantly lower CE levels in A2780 LDs grown in the presence of palmitic acid (PA), which resulted in a more balanced CE:TAG ratio (**Fig. 3B, C**). As expected, the addition of cholesterol drastically shifted LD levels and ratios towards CE, whereas subjecting cells to starvation did not significantly alter CE:TAG ratios (**Fig. 3B, C**). Collectively, these results confirmed that the native composition of LDs in A2780 cells was inherently biased towards high CE contents, which may explain their excellent suitability for αSyn binding studies. Based on these findings, we asked whether changes in CE:TAG ratios translated into observable differences in αSyn LD-binding in growth condition-challenged cells, for which we imaged endogenous αSyn in the presence of OA, PA, cholesterol and upon starvation. In line with our model and supported by the measured CE:TAG ratios, we observed uniform αSyn-LD colocalization under starvation (**Fig. 3D, Suppl. Fig. 6E**) and in the presence of OA and cholesterol (**Fig. 3D, Suppl. Fig. 6B, D**), whereas PA-treated cells exhibited greatly reduced and selective interaction patterns, with many LDs that did not colocalize with αSyn, while neighboring organelles were prominently decorated (**Fig. 3D, Suppl. Fig. 6C**). To further substantiate αSyn’s binding preference for CE-rich LDs, we took advantage of modified yeast strains that harbor deletions in either their CE (*are1Δ, are2Δ*) or TAG (*dga1Δ, lro1Δ*) biosynthesis genes ^65^. Accordingly, LDs purified from these strains were highly enriched in TAG or CE, respectively (**Fig. 3E**). We incubated both types of LDs with AF488-labeled αSyn and detected binding to CE-but not to TAG-rich LDs. Encouraged by these results, we asked whether αSyn’s binding selectivity extended to reconstituted droplet emulsions that we prepared from cholesteryl-oleate (CO) or triolein (TG) containing fluorescent nitrobenzoxadiazole (NBD)-labeled TG ^66, 67^ (**Fig. 3F**). We incubated mixtures of these droplets with AF488-labeled αSyn and found that the protein predominantly interacted with cholesteryl-oleate samples, whereas we only detected αSyn background fluorescence on NBD-triolein droplets. We obtained similar results with lipid-monolayer (PL) surrounded droplet samples (**Fig. 3F**), which emphasized the notion that cholesterol-sensing constituted a core aspect of αSyn-LD interactions. To compare the strength of αSyn binding to cholesteryl droplets with known interaction partners such as amphipathic helix (AH) peptides derived from caveolin 1 (Cav1) ^66^, we set up competition assays with pre-bound NBD-tagged AH-Cav1 droplets to which we added AF594-labeled αSyn (**Fig. 3G**). Strikingly, we observed fast and exhaustive displacement of Cav1 by αSyn, which underscored the strong avidity with which αSyn interacted with cholesteryl-oleate droplets.

Having determined that CE levels and higher CE to TAG ratios were critically important for αSyn binding to LDs (**Fig. 3**), we asked whether changing these ratios via modulations of cellular metabolic activities was capable of weakening or strengthening αSyn-LD interactions. We reasoned that starvation offered an attractive route to studying these effects, given that metabolic switching to *β*-oxidation involves multiple well-defined steps of LD processing that progressively deplete TAG and concomitantly increase CE contents ^68^, ultimately priming organelles for degradation through lipophagy ^69^. In the course of *β*-oxidation, LDs and mitochondria form extended contact sites to enable the channeling of fatty acids generated by TAG breakdown ^70^. One key enzyme in this cascade is adipose triglyceride lipase (ATGL), which catalyzes the conversion of triglycerides to diglycerides ^71^. Thus, ATGL identifies TAG-rich LDs under different metabolic conditions. Upon TAG depletion through *β*-oxidation, ATGL is removed from LDs via proteasomal degradation, which initiates organelle turnover via lipophagy ^69^. To investigate intracellular localization of αSyn during this process, we imaged LDs, mitochondria, ATGL and αSyn in A2780 cells upon glucose and serum starvation. Under nutrient-rich conditions (CM), we found mitochondria dispersed throughout the cytoplasm, whereas αSyn and ATGL colocalized on LDs (**Fig. 4A, Suppl. Fig. 7A**), as expected for the mixed TAG-CE contents we had determined earlier (**Fig. 3B**). After 24 h of starvation, LDs formed prominent ring-like membrane contact-sites (MCS) with mitochondria, indicative of active *β*-oxidation ^72^. We detected both αSyn and ATGL on LDs at this timepoint, suggesting that mitochondria-LD contacts, TAG breakdown and fatty acid channeling did not displace αSyn (**Fig. 4B, Suppl. Fig. 7B**). After 48 h, mitochondria appeared as elongated tubular structures characteristic of starvation-induced organelle remodeling ^70, 73^ that contained embedded LDs with bound αSyn. At this stage, we did not find ATGL on LDs, which indicated that their TAG contents had been largely depleted (**Fig. 4C, Suppl. Fig. 7C**). Given that αSyn persistently decorated these LDs, we reasoned that their expectedly higher CE contents made them even better binding substrates for αSyn, in further support of the previously determined CE-selectivity of αSyn-LD interactions (**Fig. 3**).

**Figure 4:**
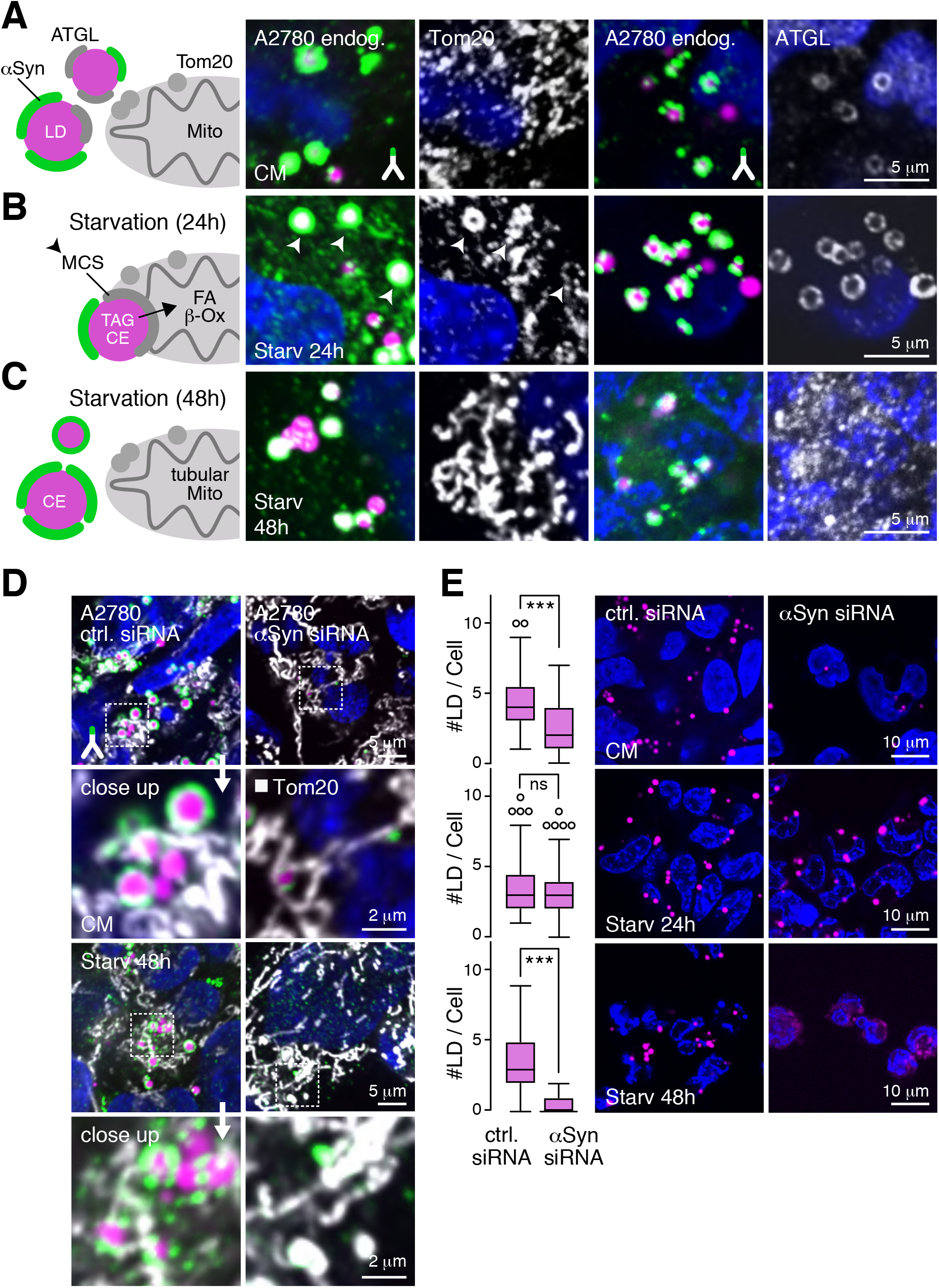
**αSyn stabilizes LDs throughout lipolysis.** (**A**) Cartoon representation of αSyn, ATGL localization on LDs and mitochondria under nutrient-rich conditions in complete medium (CM). IF-detection of endogenous αSyn (green), mitochondrial Tom20 (white) and LDs (BODIPY, purple), and of αSyn (green), ATGL (white) and LDs (purple). (**B**) Schematic of membrane contact sites (MCS) between LDs and mitochondria following glucose and serum withdrawal for 24 h, and transfer of fatty acids (FA) fueling mitochondrial *β*-oxidation (*β*-Ox). (C) Endpoint scenario of triacylglyceride (TAG) depletion and ATGL dissociation from LDs following lipolysis at late starvation time points. Representative IF images of αSyn, Tom20 and LD staining with BODIPY and of αSyn, ATGL and LDs for respective stages of starvation. (D) IF-detection of endogenous αSyn and Tom20 in A2780 cells treated with control siRNA (ctrl siRNA, left panel) and with siRNA targeting αSyn (αSyn siRNA, right panel) under nutrient-rich conditions in complete medium (CM). LDs are stained with BODIPY. Areas of close-up views are marked as dashed boxes. Bottom panels depict equivalent siRNA scenarios after 48 h of starvation. (**E**) Quantification of LD numbers per A2780 cells treated with ctrl siRNA (left row) and with αSyn siRNA (right row) at individual timepoints of starvation. Box plots (see **Materials & Methods** for details) represent 100–120 data points from three independent experiments. ns > 0.05, * p < 0.05, ** p < 0.01, *** p < 0.001 (one-way ANOVA). Corresponding images of BODIPY staining are shown on the right. For all IF images, cell nuclei/DNA are stained with DAPI (blue).

Because αSyn remained associated with LDs throughout lipolysis, we asked whether it directly contributed to this process. To answer this question, we downregulated endogenous αSyn expression with small interfering RNAs (siRNA) in A2780 cells, which we confirmed by Western blotting (**Suppl. Fig. 8A**). In complete medium and siRNA-treated cells, we noted a substantial reduction of LD numbers compared to control specimens and highly elongated mitochondria (**Fig. 4D**), similar to what we had observed earlier at late starvation stages (**Fig. 4C, Suppl. Fig. 7C**). Despite lacking LDs, some siRNA-targeted cells remained viable throughout 48 h of starvation, although their numbers were greatly reduced compared to control cells (**Suppl. Fig. 8B**). Because metabolic switching under starvation conditions involves *de novo* synthesis of LDs as an essential preparatory step for *β*-oxidation ^74^, we tested whether αSyn-deficient cells were capable of producing LDs by counting LDs in siRNA and control cells at different time points of the starvation protocol. Although siRNA cells contained significantly fewer LDs before glucose deprivation, LD numbers were comparable at 24 h of starvation (**Fig. 4E**), suggesting that αSyn-deficient cells were indeed capable of forming new LDs. However, at 48 h, pools of newly generated LDs were depleted in cells lacking αSyn, in stark contrast to native controls (**Fig. 4E**). These data suggested that loss of αSyn in A2780 cells did not impair LD biosynthesis, but affected LD turnover during starvation-induced switching to *β*-oxidation. Moreover, they insinuated that αSyn stabilized LDs and ensured their availability for lipolysis, which ultimately led to better cell survival.

Having shown that the downregulation of αSyn in A2780 cells caused a severe reduction in LD numbers even under standard growth conditions, we asked whether raising intracellular αSyn levels had the opposite effect. To address this point, we electroporated increasing amounts of αSyn into A2780 cells (**Suppl. Fig. 8C**) and counted LDs at 22 h after protein delivery. In support of our model, we measured a linear increase in LD numbers per cell, which correlated with αSyn input concentrations and plateaued at 100 μM of exogenous protein in the electroporation mixture (**Fig. 5A**). To determine whether this saturation effect was caused by the depletion of available cellular lipids needed for LD biosynthesis, we added increasing amounts of OA to electroporated cells after their recovery period of 5 h. Indeed, we measured a further increase in LD numbers that mirrored the levels of input OA (**Fig. 5A, Suppl. Fig. 8D**), with four-times as many LDs in αSyn cells compared to mock-electroporated controls in the presence of 1 mM OA. To confirm that cellular αSyn stimulated *de novo* synthesis of LDs, we probed manipulated A2780 cells with antibodies against endogenous perilipin 3 (Plin3), a marker for newly formed LDs ^75^. Whereas Plin2 colocalized with LDs in mock and αSyn-electroporated cells (**Fig. 1B, Suppl. Fig. 8E**), we only detected Plin3-decorated LDs upon αSyn delivery (**Suppl. Fig. 8E**). We also noted that LDs in mock-electroporated, OA-treated control cells were about twice as large as those in cells harboring excess αSyn (**Fig. 5B**). Considering a spherical volume relationship between LD diameters and organelle numbers, we asked whether total LD contents of αSyn and control cells were different. Surprisingly, we found that manipulated and control cells contained comparable LD volumes on a per cell basis (**Suppl. Fig. 9A**), despite significantly different organelle numbers and sizes. These observations suggested that apart from stabilizing LDs, αSyn also exerted a structural role and restricted organelle dimensions towards smaller sizes. Considering this statement in the context of our previous findings, we concluded that stabilizing and structural effects were likely functionally related, especially concerning cellular LD abundance and turnover. To test the general validity of this assumption and whether αSyn triggered the accumulation of LDs independent of whether cells contained endogenous αSyn or active LD metabolism, we turned to human embryonic kidney (HEK) cells. We did not observe LDs under standard growth conditions, in line with an earlier report ^76^, nor were we able to detect endogenous αSyn by immunofluorescence microscopy, although the LD-metabolizing enzyme ATGL was present (**Suppl. Fig. 9B**). Upon stimulation of HEK cells with OA (250 μM), we noted significant numbers of LDs that colocalized with ATGL (**Suppl. Fig. 9B**). Having verified that naïve HEK cells were capable of producing LDs, we created a stably transformed HEK cell-line that ectopically expressed wild-type αSyn under an inducible promoter. Upon induction, we confirmed progressive accumulation of intracellular αSyn by immunofluorescence microscopy and Western blotting (**Fig. 5C, Suppl. Fig. 9C**). In parallel, we counted LDs at individual time points of αSyn expression (**Fig. 5D**). In support of our model, we found that LD numbers increased linearly with αSyn levels. To determine whether direct αSyn-LD binding was required for this effect, we employed an analogous inducible HEK cell line that expressed membrane-binding deficient F4A-mutant αSyn (**Suppl. Fig. 9D**). We failed to detect LD accumulation upon F4A αSyn expression (**Fig. 5C, D**), which demonstrated that αSyn-mediated effects on LD abundance required membrane-binding. To test whether this behavior was additionally regulated by the availability of LD-forming lipids, we repeated experiments in the presence of OA (**Fig. 5E**). Despite similar overall LD numbers after 3 h of protein induction, we counted three-times as many organelles in cells expressing wild-type compared to F4A αSyn (**Fig. 5D**). Strikingly, we detected no interactions of F4A αSyn with cellular pools of LDs whereas wild-type αSyn showed pronounced organelle binding (**Fig. 5E, Suppl. Fig. 9E**). Concomitantly, LDs in F4A-expressing cells clustered in large structures of seemingly associated vesicles while organelles were dispersed in the presence of wild-type αSyn, in agreement with previous results in A2780 cells (**Fig. 5B**). In additional support of our hypothesis, these findings further corroborated that membrane-binding-competent αSyn stabilized LDs through direct interactions, which contributed to their intracellular organization and abundance.

**Figure 5:**
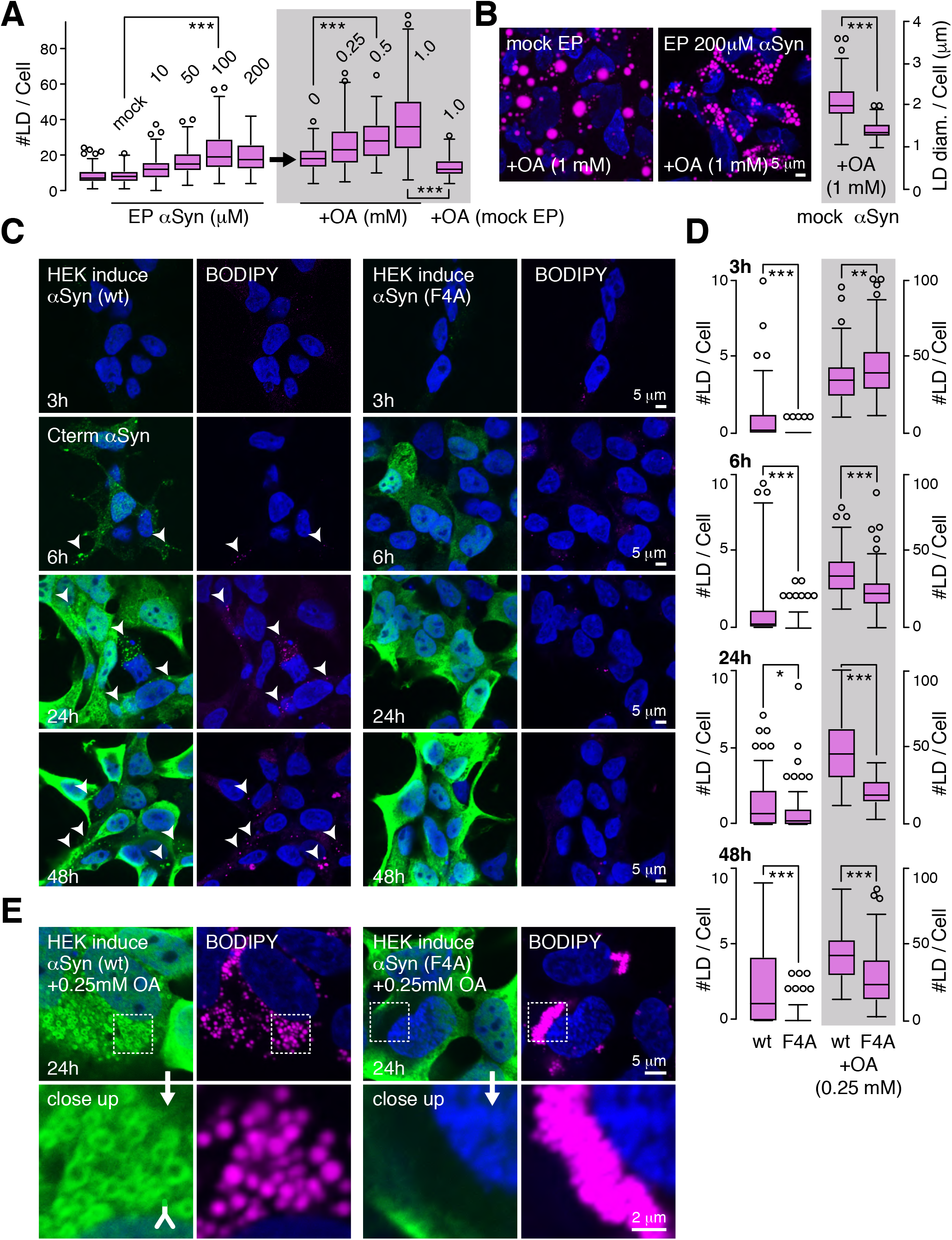
**αSyn regulates LD size and abundance.** (**A**) Quantification of LD numbers in A2780 cells following EP-delivery of increasing amounts of recombinant αSyn (10-200 μM), and for 200 μM αSyn and additional supplementation with oleic acid (OA, 0.25-1 mM). LD numbers in mock-electroporated cells in the presence of 1 mM OA are shown on the far right. Box plots represent data points from ∼120 cells collected in three independent experiments. *** p < 0.001 (one-way ANOVA). (**B**) Representative BODIPY staining of LDs in mock and 200 μM αSyn-electroporated A2780 cells in the presence of 1 mM OA and quantification of LD diameters for both conditions. Box plots represent data points (per cell) from ∼120 cells collected in three independent experiments. ns > 0.05, * p < 0.05, ** p < 0.01, *** p < 0.001 (one-way ANOVA). (**C**) IF-detection of inducible expression of wt and F4A-mutant αSyn (green) in stably transfected HEK (iHEK) cells 3 h to 48 h post induction. Corresponding BODIPY/LD staining (purple) is shown on the right. Arrowheads point to LDs upon wt αSyn expression. (**D**) Quantification of LD numbers in iHEK cells expressing wt and F4A-mutant αSyn at equivalent timepoints in the absence (left column) and presence of 0.25 mM OA (right column). Box plots represent data points from ∼100 cells, collected in two independent experiments. ns > 0.05, * p < 0.05, ** p < 0.01, *** p < 0.001 (one-way ANOVA). (**E**) Close-up images of IF-detection of wt and F4A αSyn (green) in iHEK cells 24 h post induction in the presence of 0.25 mM OA and corresponding BODIPY staining of cellular LD morphology and organization. For all IF images, cell nuclei/DNA are stained with DAPI (blue).

## Discussion

Our results demonstrate that lipid droplets in A2780, HeLa, SH-SY5Y, LUHMES, HEK, and DA-neurons from differentiated embryonic stem cells, constitute genuine intracellular binding sites for exogenously-delivered (**Fig. 1A**), transiently-overexpressed (**Suppl. Fig. 1D**), stably-integrated and induced (**Fig. 5C, 5E**), or natively-present, endogenous αSyn (**Fig. 1D**). They further establish that αSyn-LD interactions depend on membrane-anchoring properties within the N-terminus of αSyn (**Fig. 2B**) and span its first ∼100 residues (**Fig. 1C**), similar to the known binding characteristics of helical αSyn on reconstituted lipid vesicles ^15^. αSyn association and dissociation with LDs are initially dynamic but mature into more stably associated species (**Fig. 2F**), which correlates with the formation of αSyn multimers (**Fig. 2H**). We find that αSyn-LD interactions are uniquely temperature-sensitive (**Fig. 2C**) and that membrane-bound monomers/multimers are efficiently displaced below and above 37 °C (**Fig. 2D, 2E**). These results underscore the importance of temperature effects of general αSyn-membrane interactions ^77^ and may explain previous shortcomings to copurify αSyn with physiological membrane substrates such as SVs at low temperatures ^14^. It may also provide a reason for why we did not detect αSyn-LD interactions in A2780 cells in our previous experiments ^44^, which were carried out with antibodies against N-terminal αSyn epitopes and at 10 °C. Having shown that CE acts as the main determinant for αSyn binding (**Fig. 3A**) and that αSyn-LD interactions are strictly temperature sensitive within a narrow permissive range (**Fig. 2C-E**), it is likely that CE phase-transitions from isotropic liquid to ordered liquid-crystalline ^56^ translate into LD-surface properties^78^ that are more or less favorable to accommodate binding of peripheral proteins ^79^ such as αSyn ^80^ and related perilipins ^35^. In turn, cell type-specific CE:TAG ratios may determine the dynamic repertoire of LD-associated proteins ^81^, as has been observed for different members of the perilipin family ^82^, whereas changes in CE:TAG ratios, in response to TAG lipolysis for example, may induce CE phase-transitions and reshape LD proteomes, as has been demonstrated in yeast and human HeLa cells^68^. Computational models of progressive TAG depletion confirmed such CE-dependent modulations of LD-surface properties, where changes of CE:TAG ratios from 1:1 to 3:1, similar to what we measured for A2780 LDs (**Fig. 3C**), revealed greater CE interdigitation with monolayer lipids and a higher abundance of shallow lipid-packing defects ^68, 83^. Given that αSyn preferentially interacts with such defects ^80, 84^, it is likely that they contribute to the formation of productive encounter complexes between αSyn and CE-rich LDs, which may initially be weak and prone to fast dissociation, as indicated by FRAP results (**Fig. 2F**). Maturation into more stably-associated species, along with the formation of FRET-active multimers (**Fig. 2H**), hints to a second stage of LD-binding, which probably entails a deeper insertion of αSyn residues into LD cores and the corresponding masking of N-terminal epitopes (**Fig. 1C**). Considering such submerged, helical conformations along the dimensions of other monolayer-integrated helices of canonical LD proteins ^85^, αSyn may establish direct contacts with CE molecules in these states, which is supported by results from binding-assays with triolein and cholesteryl-oleate emulsions showing that αSyn can directly interact with pure cholesteryl droplets (**Fig. 3F**) and effectively displace cholesterol-binding peptides from pre-assembled emulsions (**Fig. 3G**). While we have no information about the stoichiometries and intermolecular arrangements of LD-associated αSyn multimers at this point, the extended incubation times (> 40 min) of temperature-shift experiments (**Fig. 2D** and **Fig. 2H**) suggest that they can be reversibly displaced from LD surfaces. Whether they dissociate as intact multimers or disassemble into αSyn monomers remains to be determined.

We find that endogenous αSyn-LD interactions are stably preserved throughout early and late stages of starvation-induced lipolysis (**Fig. 4A-C**). While αSyn colocalizes with mitochondrial contact sites and the LD-associated catabolic enzyme ATGL during the first 24 h of starvation (**Fig. 4B**), it remains bound to LDs after 48 h and ATGL dissociation (**Fig. 4B**), which typically coincides with TAG depletion and the formation of CE liquid-crystalline phases ^68^. We were unable to detect LD-bound αSyn undergoing fusion with lysosomes, suggesting that the protein either dissociates before the onset of lipophagy, or that C-terminal epitopes required for detection are proteolytic degraded in the process. Apart from journeying on LDs during starvation, siRNA-knockdown results indicate that endogenous αSyn stabilizes LDs in A2780 cells (**Fig. 4D**) and that the absence of αSyn leads to faster depletion of newly formed LDs without affecting their biosynthesis (**Fig. 4E**). Importantly, we find that the removal of endogenous αSyn from A2780 cells impairs their overall viability during prolonged starvation (**Suppl. Fig. 8B**), which argues for a beneficial role during lipolysis, contrary to the stipulated negative effects on TAG hydrolysis ^17^. Reciprocally, excess of intracellular αSyn increases LD numbers in dependence of lipid availability (**Fig. 5A**) and restricts LD sizes to smaller dimensions (**Fig. 5B**), which is similarly displayed in A2780 and HEK cells, with αSyn being exogenously delivered into the former and inducibly expressed in the latter (**Fig. 5C, 5D**). HEK experiments further reveal that LD-effects are strictly dependent on αSyn’s ability to interact with LD membranes and that membrane-binding-deficient F4A αSyn fails to illicit the same response (**Fig. 5C, 5D**). Thus, LD-related functions with respect to organelle accumulation and stabilization are similarly compromised upon knockdown of wild-type, or expression of F4A-mutant αSyn. This notion extends to the cellular organization of LDs, where pools of organelles appear collapsed upon F4A expression (**Fig. 5E**). Given that naïve HEK cells do not express appreciable amounts of αSyn, nor harbor LDs under standard growth conditions (**Suppl. Fig. 9B**), the correlated accumulation of LDs with induced αSyn expression corroborate reciprocal effects on LD-metabolism, which, in IMR-32 neuroblastoma cells for example, have been shown to include changes on the transcriptional and post-transcriptional level of lipid and cholesterol synthesizing enzymes and their activities ^38^. Similar LD-related changes of cellular lipid profiles haven been measured in several other cell types upon ectopic αSyn expression ^36–39^, including in non-mammalian yeast and drosophila cells ^40–42^, which, collectively, underscore the general nature of αSyn-induced effects on cellular lipid and cholesterol metabolism.

Accumulation of LDs in cells of the nervous system has received ample attention in recent years, especially in the context of ageing, cellular oxidative stress and associated lipotoxicity ^86^, and in relation to neurodegenerative disorders such as Alzheimer’s (AD) and Parkinson’s disease ^87, 88^. While healthy neurons do not accumulate LDs, they continuously require lipids and cholesterol for membrane synthesis, remodeling and homeostasis ^30^. To furnish these demands, they rely on metabolic-coupling with astrocytes, from which they take up lipid and cholesterol-laden lipoprotein particles. During ageing, cell stress, enhanced excitatory activity or upon direct fatty acid supplementation, oxidatively-damaged lipids accumulate in neurons, where they are cleared via apolipoprotein E (apoE)-mediated transport into astrocytes ^89^. Whenever lipoprotein exchange along this bidirectional axis is impaired, such as in the presence of the allelic apoE isoform 4 (apoE4), LDs accumulate in both cell types ^90^. ApoE4 constitutes the major risk gene for AD ^91^ and apoE4 astrocytes display severe defects in LD clearance ^90^, cholesterol trafficking and metabolism ^92, 93^. Similar apoE4 contributions are observed in mouse models of PD, where carriers exhibit higher amounts of insoluble αSyn aggregates, enhanced elimination of midbrain DA neurons and greater loss of postsynaptic proteins, in line with more severe disease pathology ^94, 95^. In iPSC-derived human cerebral organoids, apoE4 specimens sustain increased αSyn aggregation and pronounced accumulation of Plin2-positive, CE-rich lipid deposits, which are accompanied by reduced expression of *GBA1* and concomitant reductions in the activity of the encoded lysosomal lipid hydrolase glucocerebrosidase (GCase) ^96^. Conspicuously, *GBA1* variants are also the primary genetic risk factors for PD, whereas mutations in *GBA1* and inherited GCase deficiencies cause the lysosomal storage disorder Gaucher disease (GD), which is symptomatically associated with Lewy body dementia, a special form of PD ^97^. Most GD patients additionally develop PD and loss of GCase activity typically coincides with elevated levels of intracellular αSyn, cholesterol and CE ^98, 99^, upregulation of LD-associated genes, such as Plin4 and Plin2 ^100^, and a general accumulation of LDs ^101–103^. Thus, there is clear evidence for a common lipid-cholesterol-LD axis between the pathologies of AD, GD and PD ^99, 104^, for which αSyn’s preference for sterol-rich LDs may provide a cunning extension, especially during ageing, when lipoprotein exchange between neurons and astrocytes declines, and LDs accumulate ^86^. Under these conditions, sequestering of αSyn by LDs in neurons may induce loss-of-function effects in other parts of the cell, such as the synapse, where progressive depletion may impair vesicle trafficking and synaptic transmission. Because αSyn-binding stabilizes LDs (**Fig. 4D, E**), their accumulation may be exacerbated, probably worsening backlogging defects with astrocytes and further diminishing clearance efficiency ^89^. αSyn multimers that form on LDs (**Fig. 2G, H**) may additional contribute to such scenarios, possibly via cytotoxic gain-of-function effects similar to pathological αSyn oligomers and aggregates ^59, 60^. Whether LDs actively contribute to αSyn aggregation in the course of PD remains to be determined, although BODIPY-positive inclusions are abundantly found in *substantia nigra* neurons of PD patients ^105^ and single-layered membrane vesicles akin to LDs are discernable in brain-section micrographs of αSyn-rich Lewy body deposits ^106^. The determined sterol-selectivity of αSyn-LD interactions may indicate additionally roles in general membrane-cholesterol homeostasis, with loss-of-function defects being implicated in the clinical symptoms of many age-related dementia disorders ^107^, including PD, GD and AD ^99, 104^, but also extending to Nieman-Pick type C and Huntington’s disease ^108^. Such a cholesterol-centered framework may indeed provide answers to open questions about αSyn’s cellular functions, including its abundance in diverse sets of cells and its localization at different intracellular membrane compartments, all of which share exceptionally high cholesterol contents as their common denominator. Future investigations will clarify whether αSyn functions as such a broadly-acting, cholesterol-specific lipid-chaperone.

## Supplementary Figure Legends

**Supplementary Figure 1:**
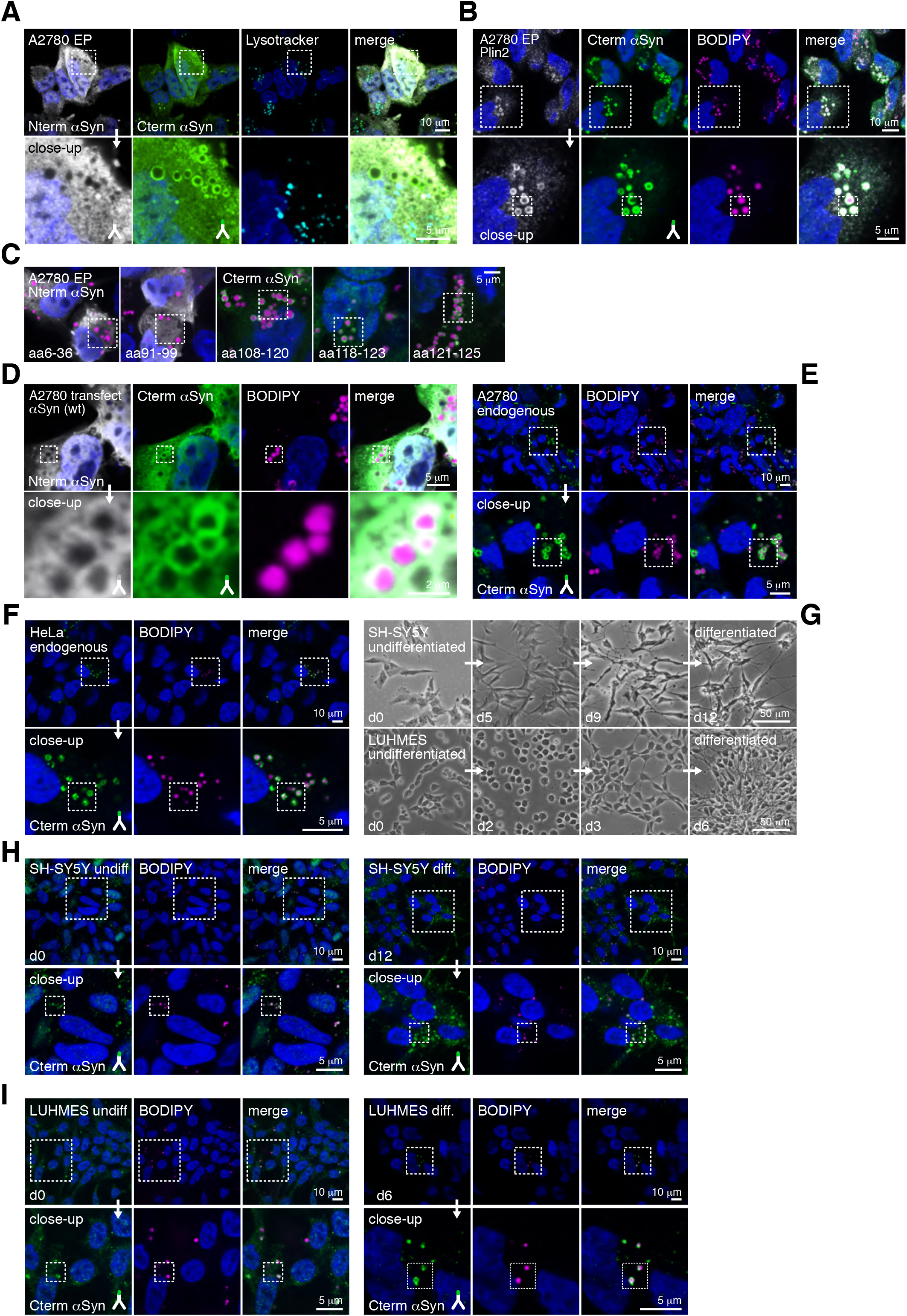
(**A**) Differential IF-detection of EP-delivered αSyn (antibodies against N- (white) and C-terminal (green) epitopes) in A2780 cells and labeling of intracellular lysosomes with lysotracker (cyan). (**B**) Low-magnification images of endogenous Plin2 (white) and EP-delivered αSyn (green) by IF-detection, and BODIPY staining of LDs (purple). Dashed boxes outline individual areas of magnification in Fig. 1B. (**C**) Low-magnification images of closeups shown in Fig. 1C. BODIPY staining (purple) identifies LDs. Dashed boxes outline magnified areas in the main figure. (**D**) Differential IF-detection of transiently transfected A2780 cells overexpressing plasmid-encoded wt αSyn. Dashed boxes outline individual areas of magnification (top to bottom). (**E**) IF-detection of endogenous αSyn in A2780 cells and BODIPY staining of intracellular LDs. Dashed boxes outline individual areas of magnification (top to bottom). Final closeup of merged αSyn and BODIPY channels is shown in Fig. 1D. (**F**) IF-detection of endogenous αSyn in HeLa cells and BODIPY staining of intracellular LDs. Dashed boxes outline individual areas of magnification (top to bottom). Final closeup of merged αSyn and BODIPY channels is shown in Fig. 1D. (**G**) Phase-contrast images of SH-SY5Y (top) and LUHMES cells (bottom) at individual days (d) of the differentiation procedures. Changes in cell morphologies indicate the progressive development of neuronal characteristics. (**H**) IF-detection of endogenous αSyn in undifferentiated (d0) and differentiated (d12) SH-SY5Y cells and BODIPY staining of intracellular LDs. Dashed boxes outline individual areas of magnification (top to bottom). Final closeup of merged αSyn and BODIPY channels is shown in Fig. 1D. (**I**) IF-detection of endogenous αSyn in undifferentiated (d0) and differentiated (d6) LUHMES cells and BODIPY staining of intracellular LDs. Dashed boxes outline individual areas of magnification (top to bottom). Final closeup of merged αSyn and BODIPY channels is shown in Fig. 1D. For all IF images, cell nuclei/DNA are stained with DAPI (blue).

**Supplementary Figure 2:**
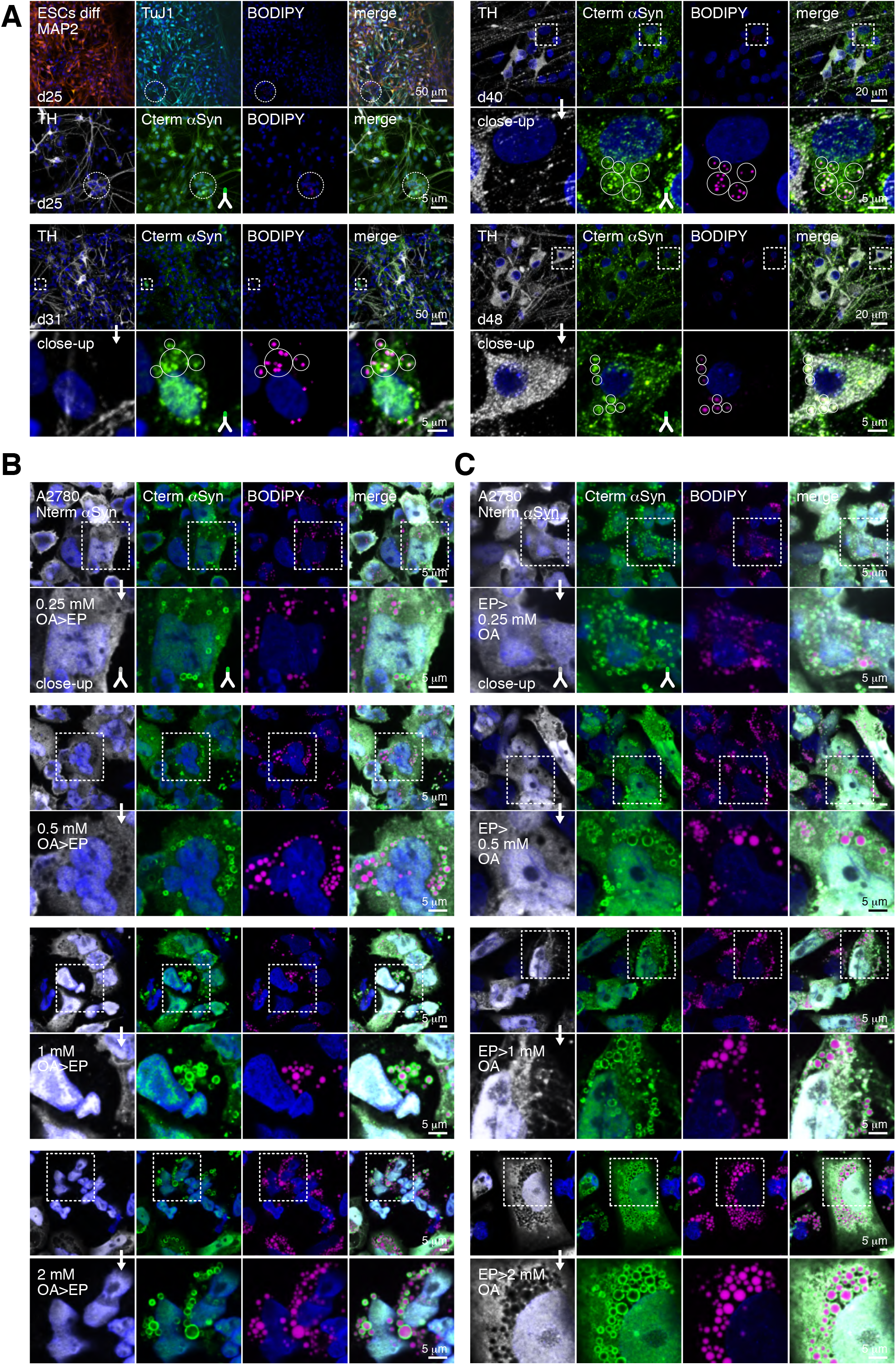
(A) Detection and intracellular localization of microtubule-associated protein 2 (MAP2), class III beta-tubulin (Tuj1) and tyrosine hydroxylase (TH), endogenous αSyn and lipid droplets (BODIPY) in human embryonic stem cells (ESCs) at day 25 (d25) of the differentiation protocol (upper left panel). TH, αSyn and LD staining at d31 (lower left panel), d40 (upper right panel) and d48 (lower right panel) of differentiation. αSyn-LD colocalization in developing dopaminergic (DA) neurons is highlighted in circles. (**B**) Differential IF-detection of N- and C-terminal αSyn epitopes in EP-transduced A2780 cells supplemented with increasing amounts of OA (0.25 mM to 2 mM, top to bottom) prior to protein delivery. Dashed boxes outline individual areas of magnification. (**C**) Differential IF-detection of N- and C-terminal αSyn epitopes in EP-transduced A2780 cells supplemented with increasing amounts of OA (0.25 mM to 2 mM, top to bottom) after protein delivery. Dashed boxes outline individual areas of magnification. For all IF images, cell nuclei/DNA are stained with DAPI (blue).

**Supplementary Figure 3:**
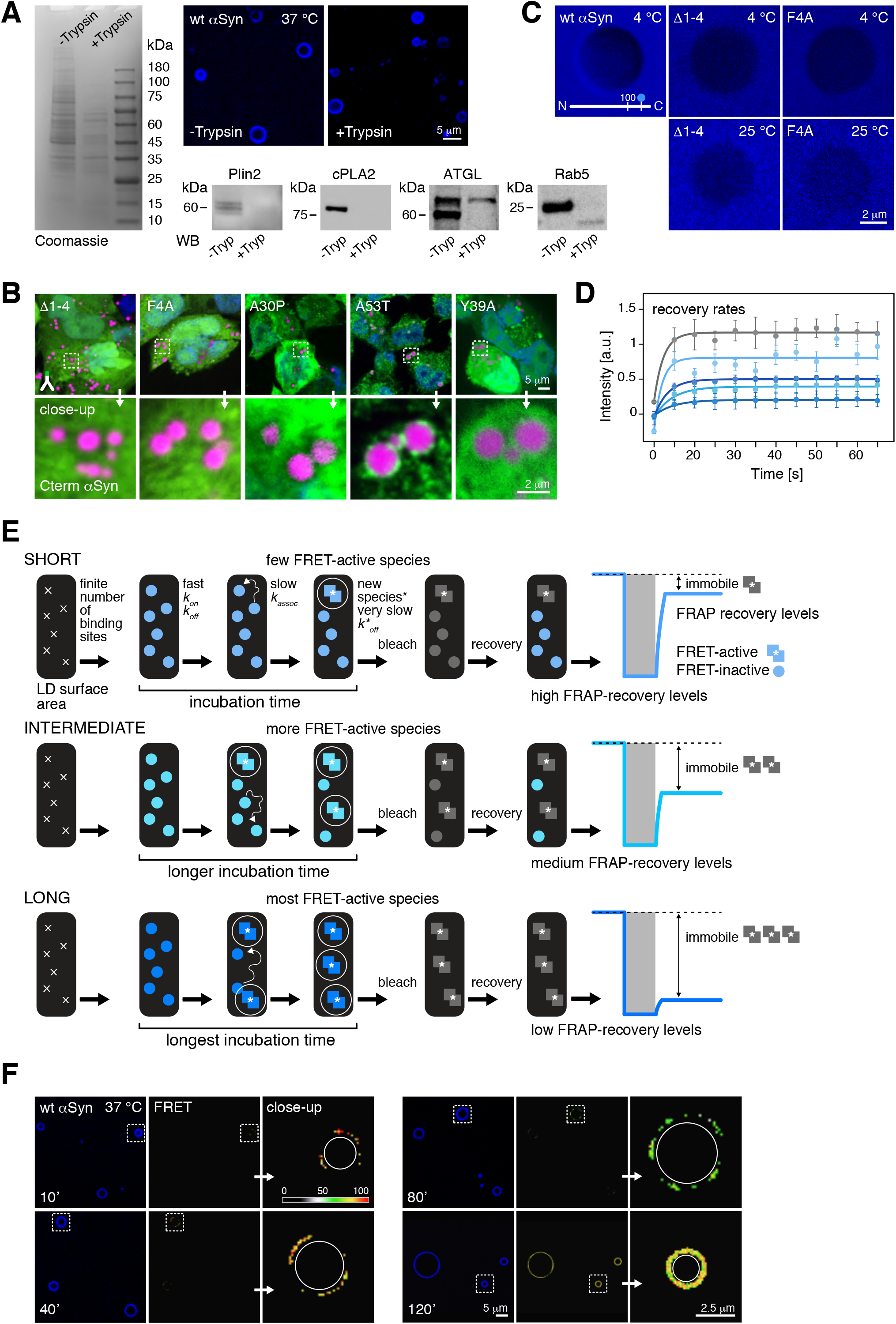
(**A**) Coomassie-stained SDS PAGE of protein components of purified LDs before (left lane) and after trypsin digestion (right lane). Western blotting (WB) of LD proteins in trypsin-treated and untreated LD samples. Fluorescence microscopy imaging of Alexa Fluor 488 (pseudo-colored in blue) labeled αSyn (N122C) binding to trypsin-treated and untreated LDs. (**B**) IF-detection of EP-delivered mutant αSyn (as shown in Fig. 2B, green) in A2780 cells. LDs are stained with BODIPY (purple), dashed boxes outline individual areas of magnification (top to bottom). Cell nuclei/DNA are stained with DAPI (blue). (**C**) Absence of binding of wt and mutant fluorescently labeled αSyn to purified LDs at 4 °C (top row) and at 25 °C (Fig. 2C and bottom row). (**D**) Fitted time-evolution of FRAP signals of αSyn in buffer (grey) and on LDs after 10 (light blue), 40 (dark blue), 80 (cyan) and 120 min (azure) of incubation at 37 °C. Individual recovery rates are estimated based on the fitted slopes connecting the first three measurement points (0, 10 and 20 s). (**E**) Schematic model outlining a possible scenario to correlate the time-evolution of FRAP and FRET results. Top row: Short incubation times of LDs and fluorescently labeled αSyn produce few FRET-active species of αSyn multimers that exhibit reduced dissociation properties. These may not be replenished via lateral diffusion of LD-bound αSyn monomers, nor with fluorescently labeled molecules in the surrounding bulk. Accordingly, recovered fluorescence is diminished by the fraction of immobile, non-exchangeable αSyn. Middle row: Intermediate incubation times increase the fraction of FRET-active, non-exchangeable αSyn species and, concomitantly, reduce respective fluorescence levels after FRAP. Bottom row: Extended incubation times produce maximum amounts of immobile αSyn multimers and lowest fluorescence recovery. While the timing of reduced FRAP and increased FRET signals is correlated, reduced dissociation of αSyn multimers (in our model) is a hypothetical assumption that is based on published results of the general behavior of membrane-associated αSyn oligomers. (**F**) Wide-field microscopy imaging of wt αSyn donor-fluorescence (blue panels) and corresponding FRET signals at 10, 40, 80 and 120 min of LD incubation at 37 °C. FRET-active LD species shown in the respective closeup frames are indicated by dashed boxes.

**Supplementary Figure 4:**
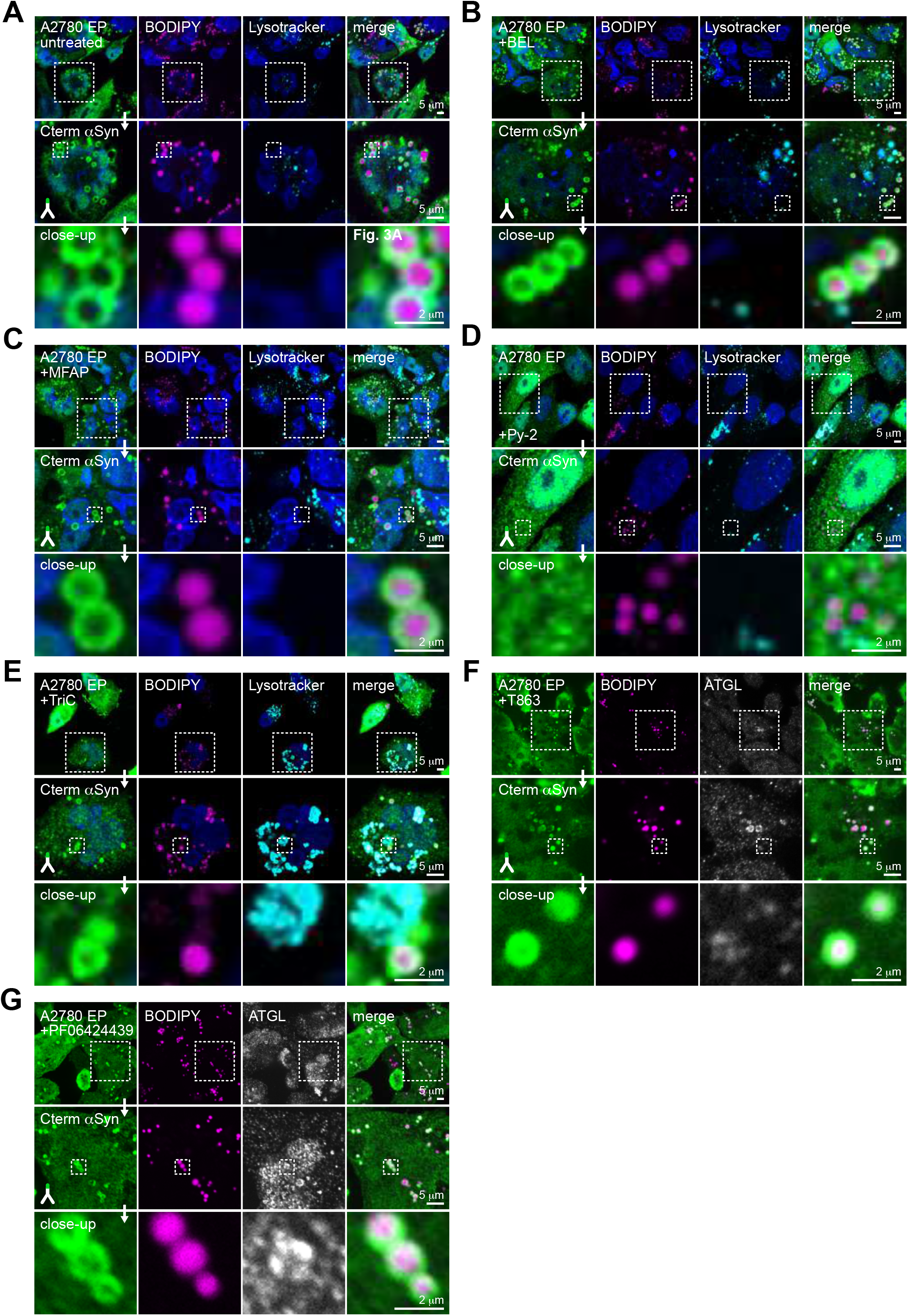
(**A**) IF-detection of EP-delivered αSyn in A2780 cells (green), colocalization with LDs (BODIPY, purple) and labeling of intracellular lysosomes with lysotracker (cyan). Low-magnification images of the merged closeup shown in Fig. 3A. Dashed boxes outline magnified areas in the main figure. (**B**) Same experimental setup, outline and coloring scheme of αSyn-transduced A2780 cells upon treatment with lysophospholipid synthesis inhibitors bromoenol lactone (BEL), (**C**) methyl-arachidonylfluorophosphonate (MAFP), (**D**) pyrrolidine-2 (Py-2) and (**E**) long chain acyl-CoA synthetase (ACSL) inhibitor Triacsin C (TriC). (**F**) IF-detection of EP-delivered αSyn and endogenous ATGL (white) in A2780 cells and colocalization with LDs, upon treatment with diacylglycerol transferase 1 (DGAT1) inhibitor T863 and (**G**) diacylglycerol transferase 2 (DGAT2) inhibitor PF-06424439. Cell nuclei/DNA are stained with DAPI (blue).

**Supplementary Figure 5:**
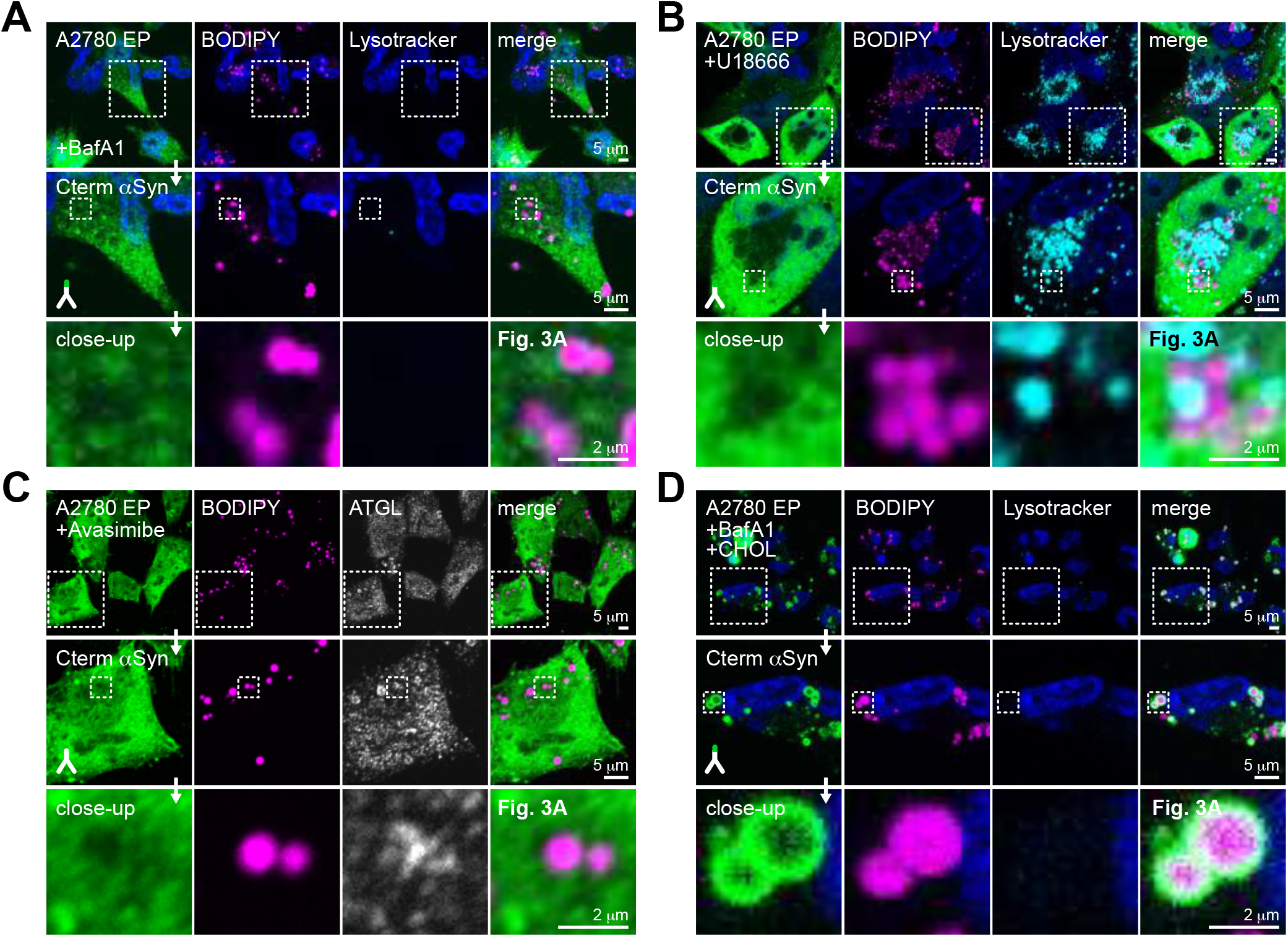
(**A**) IF-detection of EP-delivered αSyn (green) and colocalization with LDs (BODIPY, purple) in A2780 cells upon treatment with the lysosomal proton-pump inhibitor Bafilomycin A1 (BafA1). Note that labeling with the pH-sensitive dye lysotracker (cyan) does not identify lysosomes, thus confirming BafA1 activity. Low-magnification images of the merged closeup shown in Fig. 3A. Dashed boxes outline the magnified areas in the main figure. (**B**) IF-detection of EP-delivered αSyn and endogenous ATGL (white) in A2780 cells and colocalization with LDs /lysosomes (lysotracker), upon treatment with the lysosomal cholesterol transporter NPC1 inhibitor U18666 and (**C**) the CE synthetase acylCoA:cholesterol acyltransferase (ACAT) inhibitor Avasimibe. (**D**) BafA1-treated A2780 cells and rescue of αSyn-LD binding via exogenous supplementation with cholesterol. Similar to panel (**A**), absence of lysotracker staining confirms BafA1 activity. Cell nuclei/DNA are stained with DAPI (blue).

**Supplementary Figure 6:**
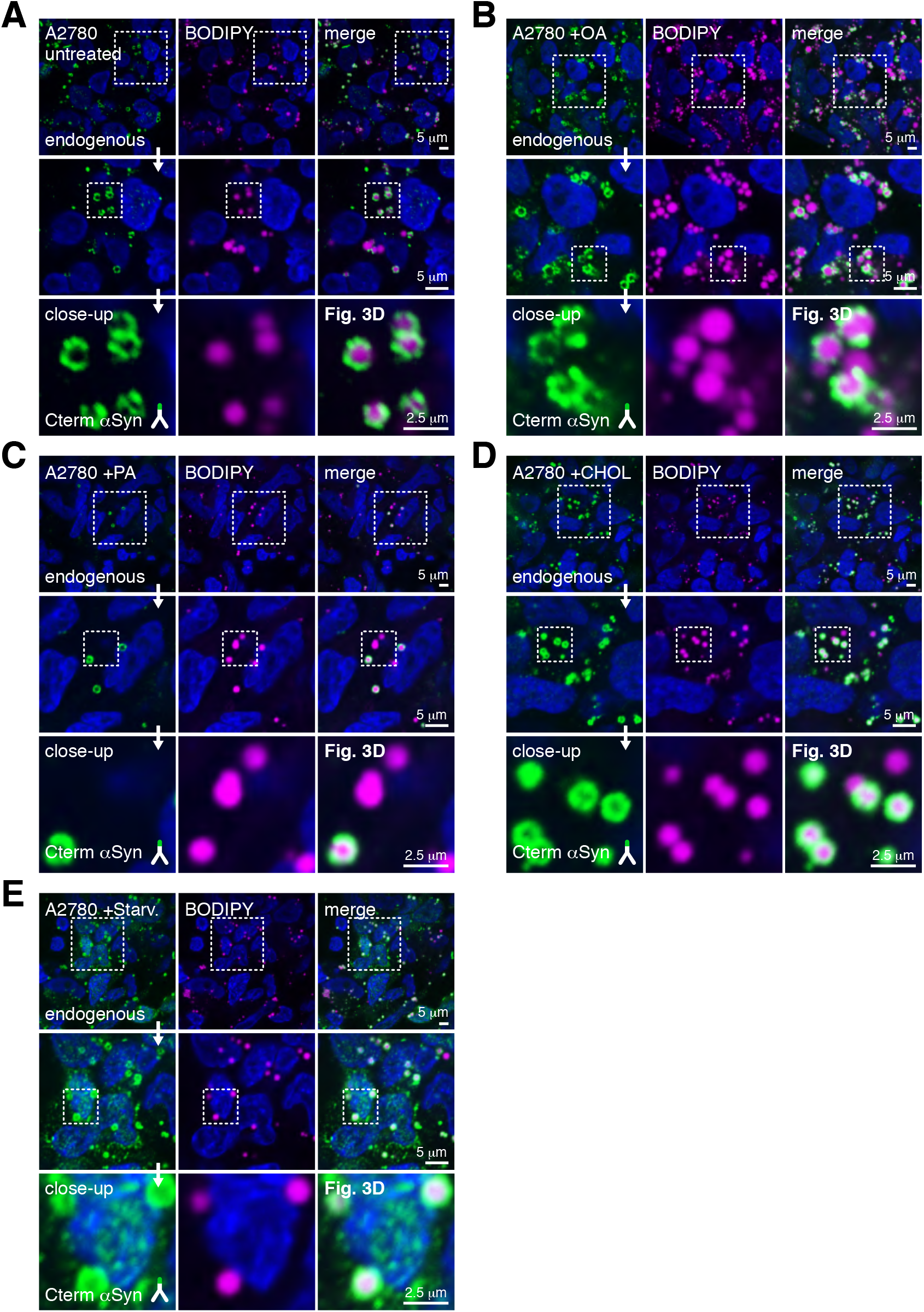
(**A**) Representative IF-detection of endogenous αSyn (green) and LDs (BODIPY, purple) in A2780 cells under complete medium conditions. (**B**) Upon supplementation with oleic acid (OA), (**C**) palmitic acid (PA), (**D**) cholesterol (CHOL), and (**E**) under starvation conditions (Starv). Low-magnification images of the merged closeup shown in Fig. 3D. Dashed boxes outline the magnified areas in the main figure. For all IF images, cell nuclei/DNA are stained with DAPI (blue).

**Supplementary Figure 7:**
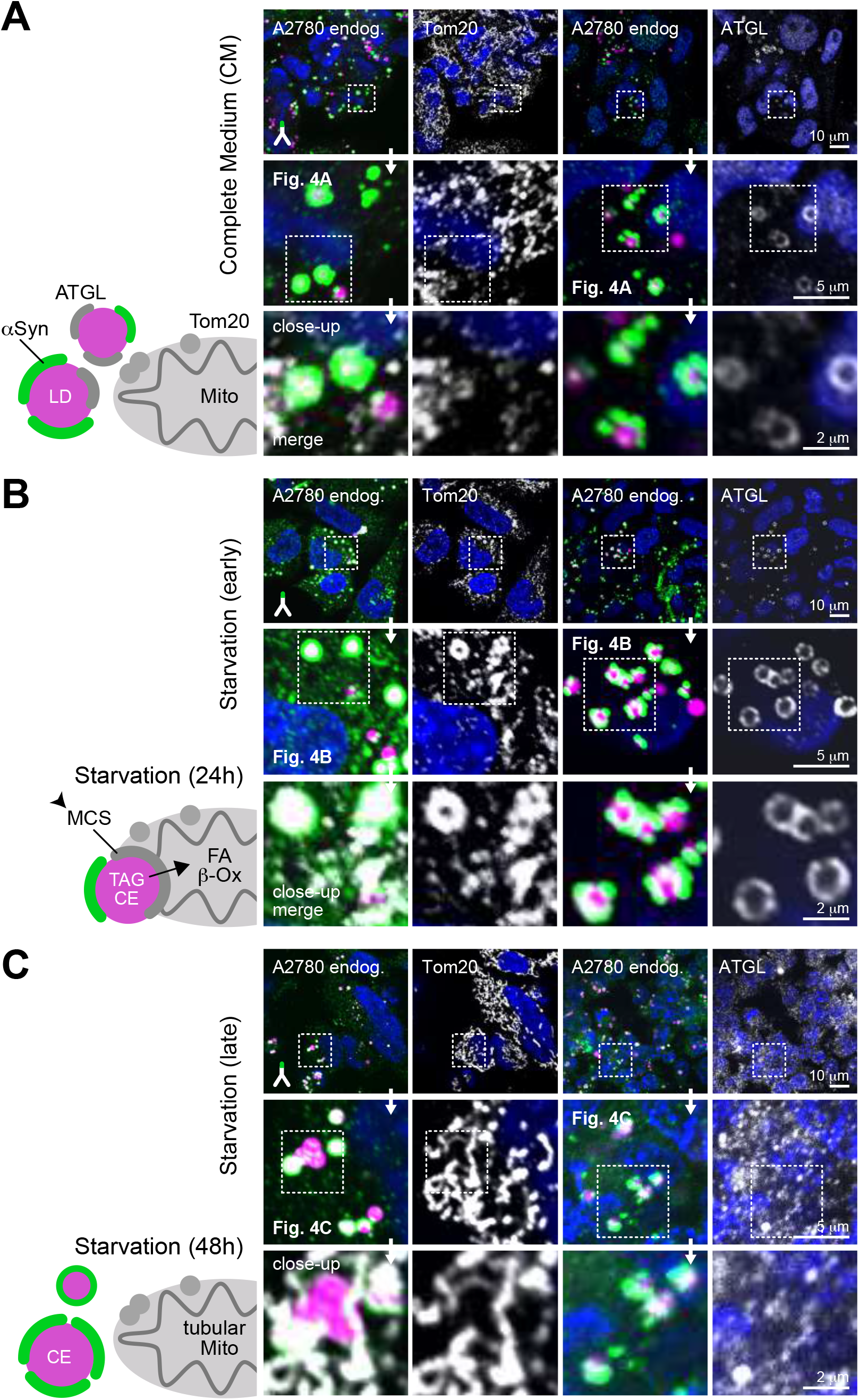
(**A, B, C**) Representative low and high magnification areas of IF-microscopy images shown in Fig. 4A, **4B** and **4C**. Dashed boxes indicate respective image sections in the closeup views. For all IF images, cell nuclei/DNA are stained with DAPI (blue).

**Supplementary Figure 8:**
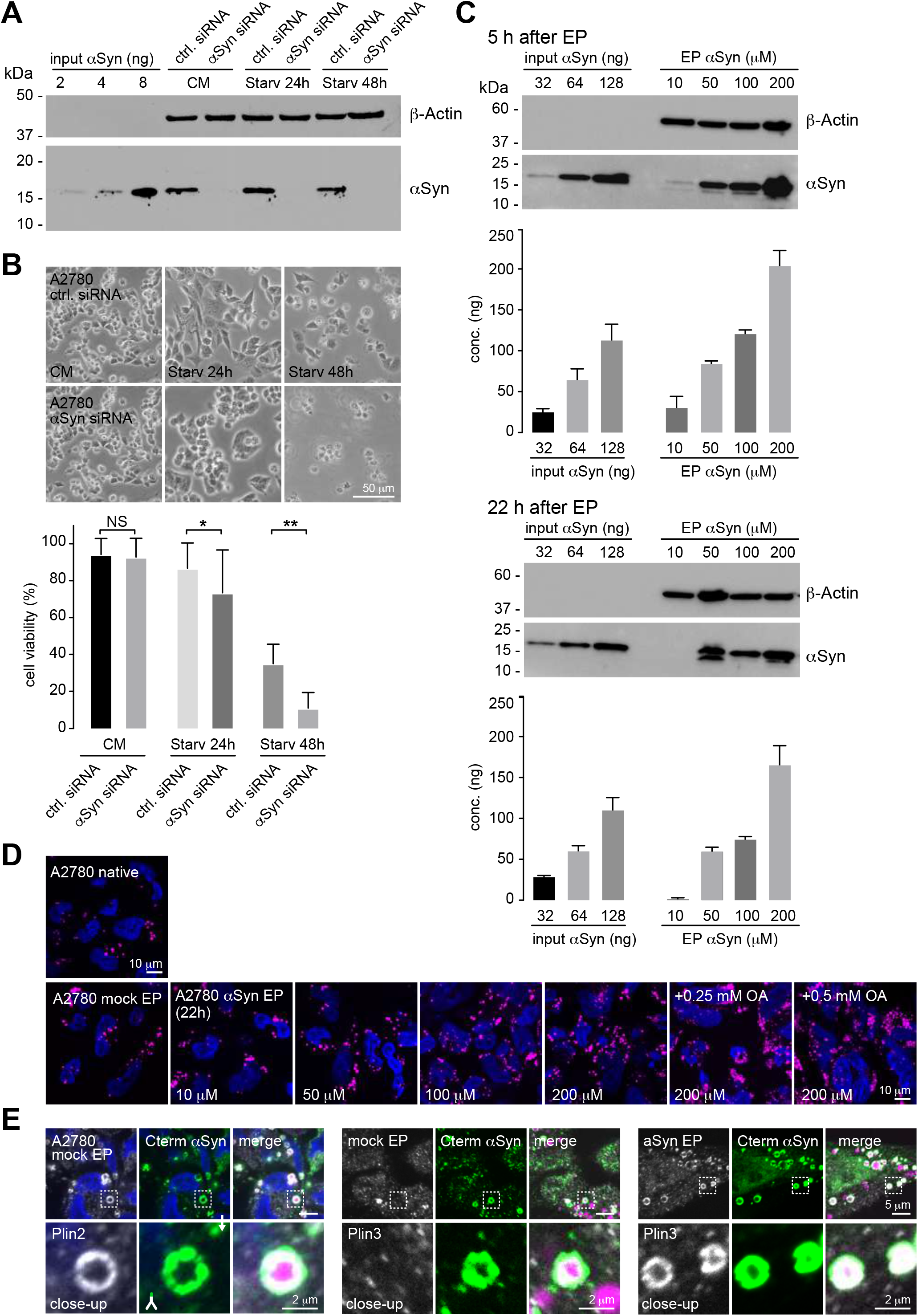
(**A**) Lysate WB detection of endogenous αSyn in control (ctrl) and αSyn-targeted siRNAs (αSyn siRNA) treated A2780 cells grown under nutrient-rich conditions (CM) and after 24 h and 48 h of starvation. 2-8 ng of recombinant αSyn serves as input control, detection of *β*-Actin signals confirms equal lysate loading. (**B**) Phase-contrast microscopy images of A2780 cells treated with ctrl and αSyn siRNA at individual starvation time-points. Cell viability as determined by Trypan blue exclusion assays and live-cell counting. Bar graphs denote percentages of cell viability and represent mean ± SD from three independent experiments. ns > 0.05, * p < 0.05, ** p < 0.01, *** p < 0.001 (one-way ANOVA). (**C**) WB quantification of intracellular αSyn concentrations in lysates of A2780 cells (50 μg per lane) 5 h and 22 h after EP delivery of the indicated molar concentrations of recombinant αSyn in the respective EP mixtures. *β*-Actin signals serve as controls for equal lysate loading. Bar graphs denote αSyn concentrations as mean ± SD from two independent experiments. (**D**) Representative images of BODIPY/LD staining (purple) of native and mock-electroporated A2780 cells and of cells harboring EP-delivered αSyn at the indicated concentrations of EP mixtures, 22 h post transduction. Panels on the right depict LD populations in A2780 cells after delivery of 200 μM αSyn and upon supplementation with 0.25 mM and 0.5 mM OA. (**E**) IF-detection of endogenous Plin2, Plin3 (white), αSyn (green) and LDs (BODIPY, purple) in mock electroporated A2780 cells (left and center panels) and of Plin3 (white) and αSyn (green), LDs (purple) in A2780 cells upon EP-delivery of 200 mM αSyn (right panels). Dashed boxes indicate individual magnification areas. Cell nuclei/DNA in all panels are stained with DAPI (blue).

**Supplementary Figure 9:**
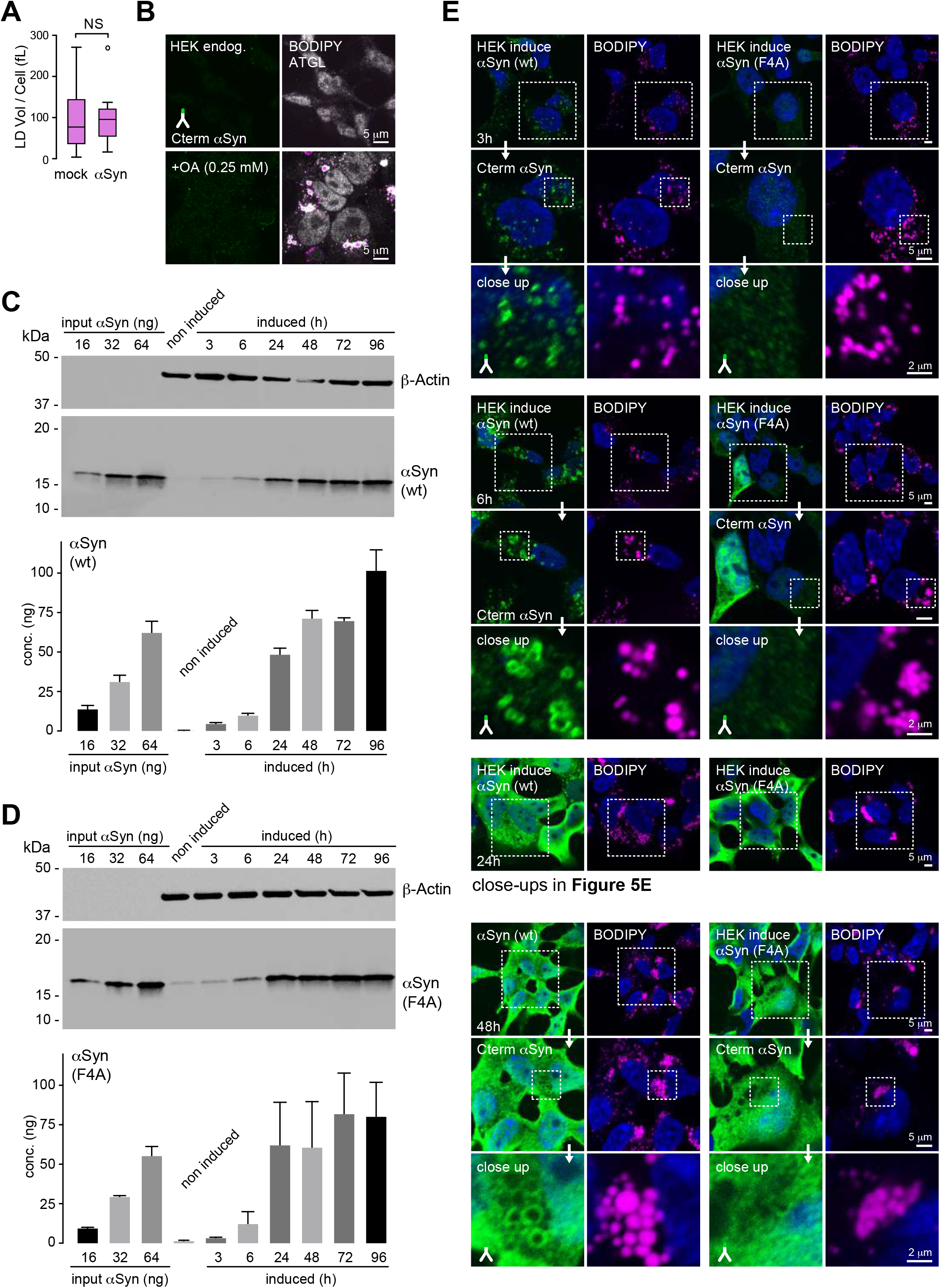
(**A**) Quantification of LD volumes of mock and αSyn-electroporated (200 μM) A2780 cells in the presence of 1 mM OA, corresponding to results shown in Fig. 5B. Box plots represent data collected per cell (n ∼ 20) from three independent experiments. ns > 0.05 (one-way ANOVA). (**B**) IF-detection of endogenous αSyn (green) and ATGL (white) in naïve HEK cells and corresponding LD levels (BODIPY, purple) in the absence (top row) and presence of 0.25 mM OA in the growth medium (bottom row). **(C)** WB quantification of intracellular αSyn concentrations in lysates (30 μg per lane) of non-induced and doxycycline (1 μg/mL) induced wt αSyn and (**D**) F4A-mutant expressing HEK (iHEK) cells 3 h to 96 h post induction. *β*-Actin signals serve as controls for equal lysate loading. Bar graphs denote αSyn concentrations as mean ± SD from two independent biological replicates. (**E**) IF-detection of wt and F4A αSyn (green) in iHEK cells 3 h, 6 h, 24 h and 48 h post induction (top to bottom) in the presence of 0.25 mM OA and BODIPY staining of corresponding LD morphologies and organizations (purple). Dashed boxes outline individual areas of magnification. Cell nuclei/DNA are stained with DAPI (blue).

## Materials and Methods

### Mammalian Cell Lines and Growth Media

Cell lines used in this study are summarized in **Table 1**. Cells were grown in humidified 5 % (v/v) CO_2_ incubators at 37 °C in the following media supplemented with 10 % (v/v) fetal bovine serum (FBS): RPMI 1640 (A2780), low glucose DMEM (HeLa,), DMEM-Ham’s F-12 (SH-SY5Y, LUHMES) and high glucose DMEM (HEK). Details for H9 human embryonic stem cells (ESCs) are outlined separately below. Cells were split at 70-80 % confluence with a passage number below 20 for all experiments. Cell lines were confirmed to be mycoplasma-free every 2 weeks.

**Table 1:**
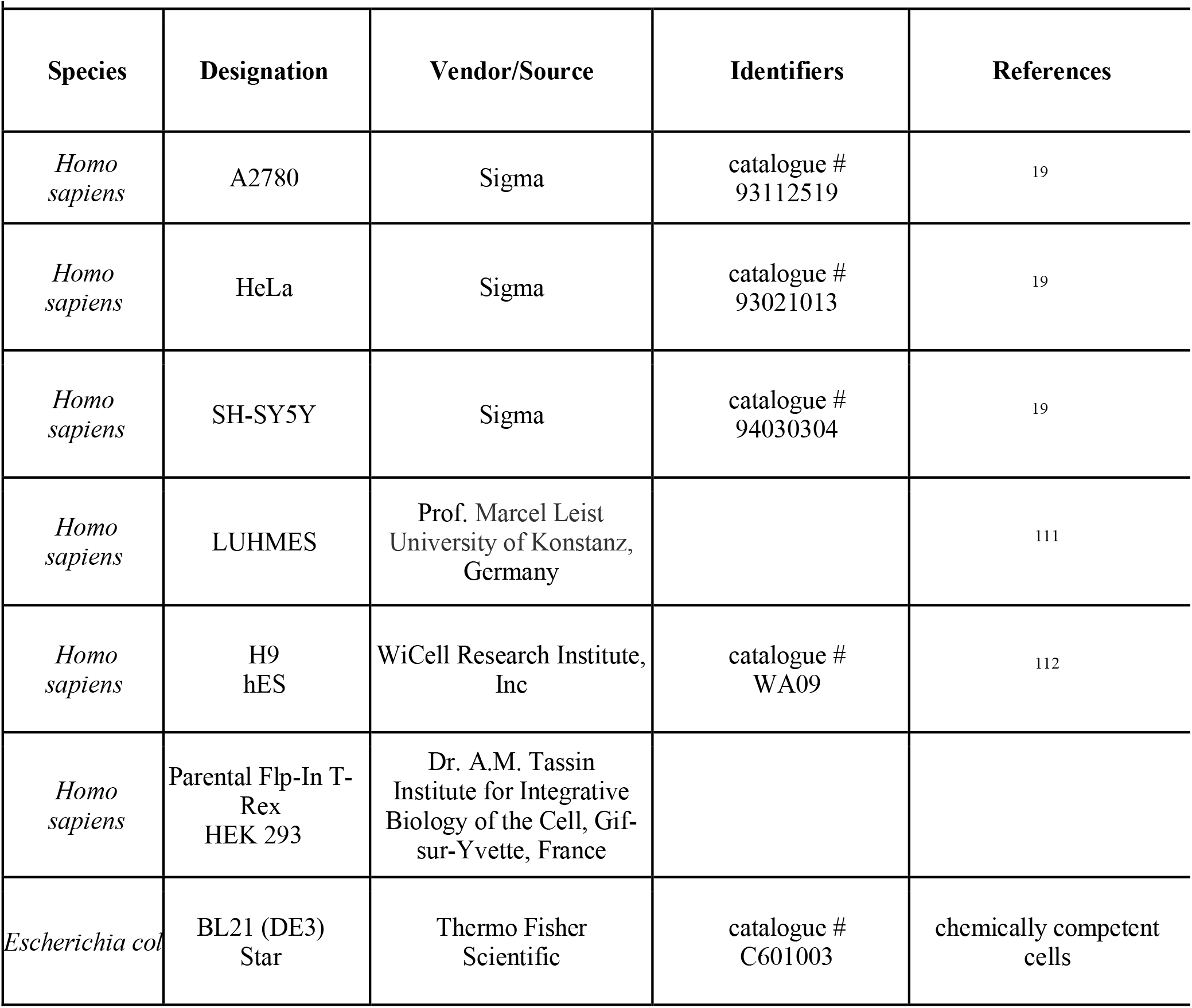
Cell Lines.

### Transient Transfection

A2780 cells were transiently transfected as described ^19^. 3 × 10^5^ cells were seeded on fibronectin (Sigma #F1141)-coated 18 mm coverslips in 12-well plates and transfected with the pHLsec-αSyn plasmid (kind gift by Radu Aricescu, University of Oxford, UK) encoding codon-optimized αSyn for optimal expression in mammalian cells using Lipofectamine^TM^ 3000 (ThermoScientific #L3000015) according to the manufacturer’s instructions. 1 µg of αSyn plasmids were used for each reaction. Following transfection, cells were grown for 24 h, washed 3 x 5 min with PBS (Gibco #14200-067) and fixed with 4 % (w/v) paraformaldehyde (PFA, ThermoScientific #28908) for 15 min at room temperature (RT) and processed for immunofluorescence (IF) microscopy. [**Fig1B, suppl. Fig1B, 1C**]

### SH-SY5Y Differentiation

SH-SY5Y cells were differentiated as described ^109^. Undifferentiated (undiff) cells were seeded at a density of 2 × 10^5^ on collagen-coated 6-well plates. 24 h after seeding, cells were treated with 10 µM all-trans retinoic acid (RA) (Sigma #R2625) in complete medium (CM). After 5 days, cells were supplemented with 50 ng/mL brain-derived neurotrophic factor (BDNF, Peprotech # 450-02) in CM without FBS and grown for another 7 days. Neuronal morphology of differentiated (diff) SH-SY5Y cells was confirmed by phase-contrast imaging on a Nikon Eclipse TS 100 microscope and cells were fixed for IF analysis. [**Fig1C, suppl. Fig1F, 1G**]

### LUHMES Differentiation

LUHMES cells were cultured and differentiated as described ^110, 111^. Undifferentiated (undiff) cells were seeded at a density of 0.5 × 10^6^ on 6-well plates precoated with 50 μg/mL poly-L-ornithine (Sigma #P-3655) and 1 µg/mL fibronectin in DMEM F12 with 1 % N-2 supplement (Invitrogen #17502048), 1 % L-Glutamine (Biological Industries #03-020-1B), 0.04 µg/mL fibroblast growth factor (FGF, R&D Systems #4114-TC) and harvested after 3 days. For differentiation, 1 × 10^6^ cells were seeded onto precoated 6-well plates in DMEM F12 with 1 % N-2 supplement, 1 % L-Glutamine, 1 µg/mL tetracycline (Sigma #T-7660), 2 ng/mL glial cell-derived neurotrophic factor (GDNF, R&D Systems #212-GD), 500 µg/mL dibutyryl cAMP (Sigma #D0627). At day 6, neuronal morphology of differentiated (diff) LUHMES cells was confirmed by phase-contrast imaging on a Nikon Eclipse TS 100 microscope and cells were fixed for IF analysis. [**Fig1C, suppl. Fig1F, 1H**]

### Embryonic Stem Cell (ESC) Differentiation

Undifferentiated H9 human embryonic stem cells (hESC, WiCell, Madison, WI) were cultured on feeder layers of irradiated mouse embryonic fibroblast (iMEFs, Stem Cell Core, WIS) in hESb medium composed of Dulbecco′s Modified Eagle′s Medium/Nutrient Mixture F-12 Ham (DMEM/F12, Sigma #D8437), 20 % KO serum replacement (KOSR, Gibco #10828028), 2 mM GlutaMax^TM^ (Gibco #35050-038), 1x MEM non-essential amino acids solution (NEAA, Sartorius #01-340-1B), 100 μM β-mercaptoethanol (β-ME, Gibco #21985023), 8 ng/mL fibroblast growth factor-2 (FGF-2, Peprotech #100-18B). Cells were passaged with collagenase IV (Worthington #LS004180). Prior to differentiation, iMEFs were removed by transferring cells to GelTrex (Gibco #A1413302) coated plates and cultured in mTeSR medium (STEMCELL Technologies #85850) for one passage. hESC were differentiated using the floorplate (FP)-derived midbrain DA neuron induction protocol as previously described ^112^ with some modifications. hESCs depleted of iMEFs were detached using StemPro Accutase (Gibco #A1110501), then seeded onto GelTrex-coated surfaces at a density of 1 x 10^5^ cells/cm^2^ and grown in mTeSR medium supplemented with 10 μM Rho-Kinase (ROCK) inhibitor (Y27632, Axon Medchem #Axon1683) for 1-2 days until 90 % confluency. At the desired confluency (day 0), mTeSR medium was replaced with KSR medium containing DMEM (Gibco #11965092), 15 % KOSR, 2 mM GlutaMax and 100 μM β-ME, along with SMAD inhibitors, i.e., 100 nM LDN 193189 (LDN, Axon Medchem #Axon1509) and 10 μM SB431542 (Axon Medchem #Axon1661). Cells were cultured for 5 days. From day 1 to 5, cells were additionally treated with 100 ng/mL sonic hedgehog (SHH) agonists SHH C24II (R&D #1845-SH), 2 µM Purmorphamine (Pur, Axon Medchem #Axon1690), 100 ng/mL Fibroblast growth factor 8 (FGF8, Peprotech #100-25A) to induce SHH signaling. From day 3 onward, differentiation medium was also supplemented with 3 µM CHIR99021 (CHIR, Peprotech #2520691) to activate wingless-related integration site (Wnt) signaling. Starting from day 5 of differentiation, SB431542 was withdrawn and KSR medium was incrementally supplemented with N2 media (from 25 %, to 50 % and 75 % over 6 days, for 2 days each) containing DMEM/F-12, 1x N-2 supplement (Gibco #17502048), 2 mM GlutaMax, 1x NEAA, 100 μM β-ME. Concentrations of LDN, SHHC24II, Pur, and CHIR remained unchanged. On day 11, the medium was switched to Neuronal Induction (NI) medium containing Neurobasal medium (Gibco #21103049), 2 % B27 supplement (Gibco #A3582801) and 2 mM GlutaMax) with 3 μM CHIR until day 13. From day 11 to 19, NI medium was further supplemented with 20 ng/mL BDNF (Peprotech #450-02), 20 ng/mL GDNF (Peprotech #450-10), 1 ng/ mL transforming growth factor type β3 (TGFβ3, Peprotech #100-36E), 10 μM DAPT (Axon Medchem #Axon1484), 0.2 mM Ascorbic Acid 2-Phosphate (Sigma #A8960) and 1 mM dibutyryl cAMP (dbcAMP, Sigma #D0627), i.e., the neuronal differentiation medium. At day 20 of differentiation, cells were dissociated using StemPro Accutase and reseeded in 8 well µ-Slides (IBDI #80826) pre-coated with 15 µg/mL poly-L-ornithine (Sigma #P-3655), 1 µg/mL laminin (Sigma #L4544), 2 µg/mL fibronectin (Sigma #F1141) at a desnsity of 35 × 10^4^ cells/cm^2^. Cells were maintained in neuronal differentiation medium and medium was exchanged every second day until fully differentiated DA neurons were formed (∼day 48). For IF analysis, cells were fixed in 8 % w/v PFA and 30 % w/v sucrose in PBS for 30 min at RT, at the indicated stages/days of differentiation. [**Fig1C, suppl. Fig2A**]

### Doxycycline-inducible HEK (iHEK) cells

Wild-type αSyn cDNA was optimized for expression in human cells and synthesized by Genscript before being cloned into a pcDNA5/FRT/TO/EGFP vector (kind gift from A.M. Tassin, Institute for Integrative Biology of the Cell, Gif-sur-Yvette, France) via BamHI/NotI restriction sites. The coding sequence for EGFP was removed by introducing a STOP codon after the αSyn coding sequence. αSyn F4A was created by mutagenesis from the parental wt αSyn plasmid (Genscript). Parental Flp-In T-Rex HEK 293 cells were cultivated in StableCell^TM^ DMEM-high glucose (Sigma #D0819) supplemented with GlutaMax^TM^ (Gibco #35050-038) and 10 % (v/v) FBS (Gibco #10270-106), (abbrev. as: DMEM-FBS) with 100 U/mL penicillin, 10 μg/mL streptomycin (Sartorius #03-031-1B), 10 μg/mL blasticidin S HCl (Gibco #A1113903) and 100 μg/mL zeocin^TM^ (InvivoGen #ant-zn-1). For generating cell lines stably expressing wt or F4A mutant αSyn, parental Flp-In T-Rex HEK 293 cells were seeded in 6-well plates at a density of 3 × 10^5^ cells per well, 24 h prior to transfection. Polyethylenimine (PEI) MAX^TM^ was used as the transfection agent (Polysciences #24765-1). Cells were washed once with PBS and a mixture of 1:1:2 (w/w) of pcDNA5-αSyn (4 μg): pOG44 Flp-recombinase expression vector (pOG44, Invitrogen #V600520):PEI in 2 mL of DMEM was added for 8 h. Following transfection, cells were washed and maintained in DMEM-FBS for another 24 h. Stable clones were selected in DMEM-FBS supplemented with 200 μg/mL hygromycin B (Invitrogen #10687010), 100 U/mL penicillin, 10 μg/mL, streptomycin, 10 μg/mL blasticidin S HCl and 100 μg/mL zeocin. Selection medium was changed every day for 2 days, and every 5 days until non-transformed cells detached. Stable clones (termed iHEK hereafter) were selected after 2 to 3 weeks, trypsinized and expanded into larger culture plates to reach ∼15 × 10^7^ cells (passage ∼20). Aliquots of 5-10 × 10^6^ cells were frozen and stored in liquid nitrogen. For experiments, iHEKs were grown under reduced selection conditions in DMEM-FBS with 100 μg/mL hygromycin B, 100 U/mL penicillin, 10 μg/mL streptomycin, 5 μg/mL blasticidin S HCl and 50 μg/mL zeocin (i.e. iHEK growth medium) at 37 °C, 5 % CO_2_. To induce *a*Syn expression, cells were seeded on 10 cm dishes at a density of 4 × 10^6^ cells (for Western blotting, WB) or on fibronectin-coated coverslips at a density of 1 × 10^5^ cells (for IF) 24 h prior to induction. *a*Syn production was stimulated with 1 μg/mL doxycycline (DOX, Sigma #D3072) in iHEK growth medium. At indicated times after DOX addition (3, 6, 24 and 48 h), growth media were removed, cells were washed twice with PBS and processed for either WB or IF. To verify *a*Syn expression, induced cells were directly lysed for WB. To analyze localization of expressed *a*Syn, induced cells on coverslips were fixed with 4 % PFA at RT for 15 min, washed thrice with PBS and processed for IF analysis. [**Fig5C, suppl. Fig9A, 9B**]

### Starvation

To establish starvation conditions, A2780 cells were seeded at a density of 2 × 10^5^ cells on 18 mm fibronectin-coated coverslips. Growth media were then replaced with FBS-free RPMI without glucose (Gibco #11879020) and grown for 24 h or 48 h. Cells cultured in complete growth media (CM) were used as control. After culturing for 24 or 48 h, cells seeded on coverslips were fixed with 4 % PFA and samples were processed for IF analysis. [**Fig4, suppl. Fig7**]

### Recombinant Protein Expression and Purification

N-terminally acetylated, human wild-type αSyn was produced by co-expressing PT7-7 plasmids with yeast N-acetyltransferase complex B (NatB) in *Escherichia coli* BL21 Star (DE3) cells in LB media as described ^44^. Protein purification under non-denaturing conditions was performed as published ^44^. *a*Syn mutants *Δ*N, F4A, Y39A, A30P, A53T and E46K were generated by site-directed mutagenesis (QuikChange, Agilent #200523) and confirmed by DNA sequencing. For site-specific fluorescence labeling of wild-type (wt) and mutant αSyn, Asn122 was replaced with a cysteine residue (N122C) by site-directed mutagenesis. Recombinant expression and purification of all variants of *a*Syn was identical to the wild-type protein. 5 mM β-mercaptoethanol (Sigma #m6250) was included in all purification steps of *a*Syn N122C variants to preserve the reduced state. Because *a*Syn *Δ*N lacked N-terminal consensus sites for NatB, it was produced in the non-acetylated form. Protein samples were concentrated to 1-1.2 mM in phosphate buffer (25 mM sodium phosphate, 150 mM NaCl) at pH 7.0. *a*Syn N122C variants were stored in phosphate buffer containing 10 mM Tris(2-carboxyethyl)phosphine hydrochloride (TCEP, Sigma #646547). Protein concentrations were determined spectrophotometrically by UV absorbance measurements at 280 nm with ε = 5690 M^-1^cm^-1^ for wt and all αSyn mutants. Final aliquots of protein stock solutions were snap frozen in liquid nitrogen and stored at -80 °C until use.

### Fluorophore Labeling

Site-specific fluorophore labeling of recombinant N122C αSyn was performed as described ^113^. Prior to fluorophore coupling, 500 µL aliquots of 200 µM wt or mutant αSyn were incubated with 5 mM TCEP for 30 min at RT. TCEP removal and buffer exchange into 20 mM Tris (pH 7.4), 50 mM NaCl, and 6 M guanidine hydrochloride (GdmCl) was carried out on PD10 desalting columns (GE Life Sciences #17085101). Reduced protein solutions were incubated with 2-fold molar excess of C5 maleimide Alexa Fluor 488 (AL488, Invitrogen #A10254) or Alexa Fluor 594 (AL594, Invitrogen #A10256) for 2 h (stirring) at RT. Fluorophore-coupled proteins were separated from unreacted dye using PD10 desalting column, concentrated and buffer exchanged to phosphate buffer using Amicon Ultra-4 Centrifugal Filter Unit (Millipore #UFC800324). Protein concentrations were determined spectrophotometrically by correlative absorbance measurements at 280 nm (protein), and 493 nm (absorption max AL488), 588 nm (absorption max AL594), respectively, according to:

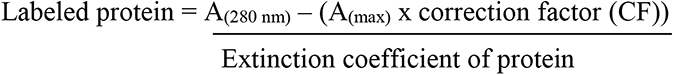

With A_(280 nm)_ as the absorbance of labeled αSyn at 280 nm, A(max) as the absorbance at the dye absorption maximum (nm) and with CF for respective dyes of CF_280_ (AL488) = 0.11; CF_280_ (AL594) = 0.56.

### Electroporation (EP)

Electroporation was performed as described ^44^. A2780 cells were detached from culture flasks with trypsin/EDTA (0.05 % / 0.02 %) (Sartorius #03-054-1B), centrifuged at 213 x g for 5 min at 25 °C, washed once in PBS and counted on an automated cell counter (BioRad, T20). Recombinant wt and mutant αSyn were added to freshly prepared and sterile filtered electroporation buffer (EPB, 100 mM sodium phosphate, 5 mM KCl, 15 mM MgCl_2_, 15 mM HEPES, 2 mM ATP (Thermo Fisher #R0441), 2 mM reduced glutathione (Sigma # G4251) at pH 7.0) to final concentrations of 10-200 µM. Pelleted cells were mixed with protein in EPB at 70 × 10^6^ cells per mL. 100 µL aliquots (7 × 10^6^ cells) were then transferred into 2 mm EP cuvettes and electroporated on an Amaxa Nucleofector I (Lonza). For A2780 cells, the B28 pulse-program was used. Cells were pulsed twice with gentle mixing in between pulses. For control experiments, cells were mock-electroporated with EPB alone. Directly after electroporation, 1 mL of pre-warmed (37 °C) CO_2_-adjusted growth medium was added to each cuvette and electroporated cells were centrifuged to remove debris. Cell pellets were resuspended in fresh media. For IF analysis, 0.5 × 10^6^ post-EP cells were seeded on 18 mm fibronectin-coated glass coverslips and cultured for 5 h under standard incubator settings (37 °C, 5 % CO_2_) for recovery. Post recovery, fresh medium was added to cells and they were grown for another 16 h. Individual coverslips were washed thrice with PBS and fixed with 4 % (w/v) PFA for 15 min at RT. Cells were washed with PBS and processed for IF analysis. Alternatively, post-EP cell pellets were seeded at a density of 4 × 10^6^ into 10 cm culture dishes and allowed to recover for 5 h. Thereafter, medium was replaced and cells were harvested by trypsinization at the indicated time points and processed for WB. [**Fig1A, suppl. Fig1A, suppl. Fig3B, suppl. Fig8C, 8D**]

### Immunofluorescence (IF) Microscopy

For lysosome staining, LysoTracker Red (ThermoScientific #L7528) was used according to manufacturer’s instructions. Briefly, A2780 cells grown on fibronectin-coated coverslips were incubated for 30 min with 75 nM LysoTracker in CM at 37 °C (in CO_2_ incubators). Following, excess dye was removed, cells were washed in pre-warmed PBS and fixed in 4 % (w/v) PFA at RT. After fixation, cells were processed for antibody staining. For IF microscopy of *a*Syn and other proteins, cells were processed as described ^19^. Briefly, after fixation and removal of PFA by washing twice by PBS, cells were permeabilized with 0.1 % Triton^TM^ X-100 (Sigma #X100) in PBS for 5 min, and blocked with 0.13 % fish skin gelatin (Sigma #G7765) in PBS for 60 min. After blocking, cells were incubated with primary antibodies for 120 min at RT. Antibody details are provided in **Table 2**. After washing thrice for 5 min with PBS, cover slips were incubated with Alexa fluorophore-tagged secondary antibodies for 60 min at RT. For LD staining, BODIPY™ 493/503 (BODIPY, ThermoScientific #D3922) was added at a concentration of 4 μM along with secondary antibodies. Before IF analysis, cover slips were counterstained with DAPI (Invitrogen #D3571) and mounted with ImmuMount (Epredia #9990402) after three PBS washes (5 min each). IF microscopy was performed on a Nikon Eclipse Ti2 microscope with a Yokogawa CSU-W1 confocal scanner unit and equipped with a Photometrics Prime 95B CMOS camera (Nikon Ti2 CSU-W1) and a Nikon CFI Plan Apochromat 100x objective with a numerical aperture (NA) of 1.4. Four channels in 7 optical sections from the central plane near the cell nucleus were acquired with excitation wavelengths of 405 nm (blue, 20 % laser power), 488 nm (green, 10 % laser power), 561 nm (red, 40 % laser power) and 631 nm (far-red, 40 % laser power) with 200 ms exposure times. At least 10 images per biological replicate were collected and 3-4 replicates per experiment were analyzed. [**Fig1, suppl. Fig1, suppl. Fig2, suppl. Fig3B, Fig3A, 3D, suppl. Fig4-7, Fig4, suppl. Fig8D, 8F, Fig5B, 5C, 5E, suppl. Fig9C**]

**Table 2:** Antibodies.

| Table 2: Antibodies |  |  |  |  |
| --- | --- | --- | --- | --- |
| Species Resource | Designation & Epitope | Vendor | Identifiers | Concentration used |
| mouse monoclonal | anti-αSyn<br>aa 121-125 | Santa Cruz | catalogue # sc69977 | IF 1:200<br>WB 1:100 |
| rabbit monoclonal | anti-αSyn<br>aa 6-36 | Abcam | catalogue # ab51252 | IF 1:200 |
| mouse monoclonal | anti-αSyn<br>aa 91-99 | Becton Dickinson | catalogue #<br>610786 | IF 1:100 |
| sheep monoclonal | anti-αSyn<br>aa 108-120 | Abcam | catalogue # ab21976 | IF 1:200 |
| rabbit monoclonal | anti-αSyn<br>(MJFR1)<br>aa 118-123 | Abcam | catalogue #<br>ab138501 | IF 1:200<br>WB 1:10000 |
| rabbit monoclonal | anti-MAP2<br>aa 1-100 | Abcam | catalogue # ab32454 | WB 1:1000 |
| mouse monoclonal | anti-Neuron<br>specific $\beta$ III<br>Tubulin | R&D Systems | catalogue #<br>MAB1195 | IF 1:1000 |
| rabbit monoclonal | anti-Tyrosine<br>Hydroxylase<br>aa 1-444 | Abcam | catalogue #<br>ab112 | IF 1:1000 |
| mouse monoclonal | anti-β-Actin | Abcam | catalogue #<br>ab6276 | WB 1:5000 |
| Guinea pig polyclonal | anti-perilipin3 (Plin3) | Progen | catalogue # GP30S | IF 1:200 |
| rabbit polyclonal | anti-perilipin2 (Plin2) | Sigma | catalogue # HPA016607 | IF 1:200 |
| rabbit polyclonal | anti-perilipin2 (Plin2)<br>aa 88-437 | Proteintech | catalogue # 5294-1AP | WB 1:1500 |
| rabbit polyclonal | anti-cytosolic Phospholipase A2 (cPLA2) | Abcam | catalogue # ab53105 | WB 1:1000 |
| rabbit monoclonal | anti-Adipose Triglyceride Lipase (ATGL)<br>aa 350-500 | Abcam | catalogue # ab220738 | IF 1:200<br>WB 1:1000 |
| rabbit monoclonal | anti-Rab5 (C8B1) | Cell Signaling Technology | catalogue # 3547 | WB 1:1000 |
| goat polyclonal | anti-mouse IgG Alexa 647 conjugated | Abcam | catalogue # ab150119 | IF 1:1000 |
| donkey polyclonal | anti-rabbit IgG Alexa 555 conjugated | Invitrogen | catalogue # A-31572 | IF 1:1000 |
| goat polyclonal | anti-mouse IgG HRP conjugated | Sigma | catalogue # A9917 | WB 1:10000 |
| goat polyclonal | anti-rabbit IgG HRP conjugated | Jackson ImmunoResearch Laboratories | catalogue # 111-035-003 | WB 1:5000 |

### Lipid and Cholesterol Treatment

Oleic acid (OA, Sigma #O1383) was dissolved in serum-free RPMI supplemented with 10 % fatty acid-free BSA to prepare 10 mM stock solutions, as published ^114, 115^. For OA treatment before EP, A2780 cells were seeded in 150 mm dishes and grown to 60 % confluence. 0.25, 0.5, 1.0 and 2.0 mM of OA was added to growth media for 24 h. Cells were harvested for EP by trypsinization as described. 200 μM αSyn was used in the respective EP mixtures. After EP, cells were seeded at a density of 0.5 × 10^6^ on fibronectin-coated 18 mm coverslips in 12-well plates and grown for 5 h. Thereafter, cells were fixed with 4 % PFA and processed for IF analysis. For treatment in the reverse order, A2780 cells were first electroporated with 200 μM αSyn and seeded at a density of 0.5 × 10^6^ on fibronectin-coated 18 mm coverslips in 12-well plates and grown for 5 h. Following, growth media were exchanged for 0.25, 0.5, 1.0 and 2.0 mM OA-supplemented ones. Cells were grown for 18 h, fixed with 4 % PFA and processed for IF analysis [**Fig1D, suppl. Fig2B, 2C, suppl. Fig8D**]. For analyzing endogenous αSyn on lipid droplets with different TAG and CE contents, A2780 cells were seeded at a density of 2 × 10^5^ on fibronectin-coated 18 mm coverslips in 12-well plates and grown for 5 h. Growth media were replaced with media supplemented with 1 mM OA, 0.5 mM palmitic acid (PA, Sigma #P0500), or 100 μM cholesterol (CHOL, Sigma #C3045). PA-conjugation was carried out with fatty acid-free BSA ^114, 115^, CHOL-conjugation was performed with methyl-*β*-cyclodextrin (M*β*CD, Sigma #C4555) ^116^. Cells were grown for 24 h, fixed with 4 % PFA and processed for IF analysis. For LD purification from different growth conditions, A2780 cells were seeded into 150 mm dishes and grown to 60 % confluence. Growth media were exchanged with media supplemented with lipid/cholesterol and cells were cultured for 24 h. Following, cells were trypsinized and collected by centrifugation at 200 × g for 5 min at 25 °C, washed once in PBS and counted on an automated cell counter (BioRad, T20). For each condition 90 × 10^6^ cells were collected and processed for lipid droplet purification [**Fig3D, suppl. Fig6**]. For OA treatment of HEK cells inducibly expressing αSyn, iHEK lines were seeded onto fibronectin-coated 18 mm coverslips at a density of 1 × 10^5^ cells 24 h prior to induction. Following, growth media were exchanged with media containing 1 μg/mL DOX, supplemented with 250 μM OA were indicated. After 3, 6, 24 and 48 h of DOX addition, media were removed and cells were washed twice with PBS, fixed with 4 % PFA and processed for IF analysis. iHEK cells cultured in growth media without DOX and supplemented with 250 μM OA were used as control. [**Fig5C, 5E, suppl. Fig8F, suppl. Fig9C**]

### Lipid Droplet (LD) Purification

LDs were purified from cell pellets as described ^117^ with some modifications. 9 × 10^6^ cells/mL cells were resuspended in PBS (Sigma #D8662) with cOmplete proteinase inhibitor cocktail (Roche #04693132001) and lysed by repeated freeze-thaw cycles. Homogenates were centrifuged at 3.000 × g for 10 min at 4 °C to remove cell nuclei, membranes and intact cells. Postnuclear supernatant (PNS) fractions were transferred into swinging-bucket centrifugation tubes (Thermo Scientific #03699), to a volume of ∼12 mL with PBS and separated by ultracentrifugation at 160.000 × g for 2 h (4 °C) in a SW Ti41 rotor (Beckman Coulter #331362). Top-layer LDs were collected by aspiration using a pre-wetted 200 µL pipette tip so that maximal LDs were collected in minimal buffer volume. LDs were resuspended in PBS (final volume ∼12 mL) and centrifuged a second time at 20.000 × g for 20 min (4 °C) for washing. After centrifugation, LDs were transferred to Eppendorf tubes and concentrated by centrifugation at 16.000 × g for 10 min (4 °C). The bottom solution was removed with a pre-wetted gel-loading tip. LD quality and concentration were assessed by phase-contrast microscopy on a Nikon Eclipse TS 100 microscope at 40x magnification. LD numbers were counted on a hemocytometer and LD-slurries containing 1 × 10^5^ LDs per μL were prepared as 100 μL aliquots, snap frozen in liquid nitrogen and stored at -80 °C until use.

### LD Interaction Assays

A2780 cells were treated with 1 mM OA and LDs were purified as outlined above. Freshly prepared or thawed LDs were mixed with fluorophore-labeled wt or mutant αSyn to final protein concentrations of 1 μM. Mixtures were incubated at 4 °C, 25 °C, or 37 °C for indicated time periods on a shaking platform. To image protein-LD interactions, LDs were transferred to home-built imaging chambers prepared with a commercial single-hole puncher (1/4”) and double-sided tape. Perforated tape was attached to a cleaned glass slide, chambers were loaded with 2 μL of LD-protein solutions and sealed with 2-

[methoxy(polyethyleneoxy)propyl]trimethoxy-silane (mPEG) coated coverslips for imaging by confocal IF microscopy. For temperature-gradient/cycle experiments, glass slides were put on temperature-adjusted thermo-blocks or on ice before imaging. Fixed-temperature time series were recorded on slides removed from thermo-blocks at indicated time points. Samples were imaged on a Nikon Ti2 CSU-W1 with a 100x objective with a NA of 1.4. For each condition, binding characteristics were independently confirmed for 2-3 batches of individually prepared LDs as biological replicates. [**Fig2A, 2B, 2C, 2H, suppl. Fig3C**]

### Fluorescence Recovery After Photobleaching (FRAP)

For FRAP experiments, AL488-labeled αSyn was incubated with purified LDs at 37 °C for the indicated time periods before transferring mixtures to imaging chambers. Slides were analyzed on an inverted Leica SP8 STED3X confocal microscope equipped with internal Hybrid (HyD) detectors, acousto-optical tunable filters (AOTF) (Leica Microsystems CMS GmbH, Germany), white light laser (WLL, excitable wavelengths 470 - 670 nm) and an 86x water immersion objective (HC PL APO 86x/1.20 W motCORR STED white). Photobleaching of αSyn signals was carried out with an Argon laser operating at 488 nm (80 % laser power, for 70 s). Six pre-bleach and 69 post-bleach images were recorded to measure fluorescence recovery every 5 s. FRAP time series were analyzed in Fiji ^118^ as described ^119^. Background and photo-bleaching effects during acquisition were corrected using the mean intensity value of FRAP zone (I_frap_), background zone (I_BG_) and total intensity (I_tot_) for each acquisition frame according to

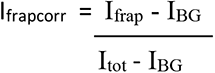

The corrected I_frap_ (I_frap corr_) was then normalized with the averaged pre-bleach intensity values of each FRAP zone (I_frap-prebleach_). Pre-bleach fluorescence was set to 1 and individual FRAP curves and recovered fluorescence intensities were determined accordingly. Mean FRAP curve data points and standard deviations were calculated from 4 technical replicates. Post-bleach FRAP data points (t = 65-120 s) were fitted to f(t) = a × (1-exp(b × t)) + c using the curve-fitting toolbox in MATLAB. Fitted coefficients had confidence intervals of 95 %. [**Fig2D, suppl. Fig3D**]

### Förster Resonance Energy Transfer (FRET)

For FRET experiments, purified LDs were mixed with fluorophore-tagged αSyn samples (wild-type or F4A mutant) that were either labeled with AL488 or AL594 (1 μM each) to enable resonance energy transfer between the two dyes. Mixtures were incubated at 37 °C for 2 h on a shaking platform. For time-resolved FRET experiments, samples were removed at indicated time points and transferred to imaging chambers. All samples were imaged on a Nikon Ti2 CSU-W1 with a 100x objective of NA 1.4 on a 37 °C heated stage mounted on an Oko lab cage incubator. FRET measurements were performed with lasers operating at 488 and 561 nm. Donor AL488 excitation was at 488 nm (laser power 5 %), with emission at 535 nm. Acceptor AL594 excitation was at 561nm (laser power 5 %), with emission at 620 nm. FRET-signals were recorded at 620 nm after excitation at 488 nm (power 5 %) for 200 ms (exposure time). Mixtures of LDs and AL488 αSyn, or LDs and AL594 αSyn, were first imaged separately in FRET mode to determine background (BG) intensities and spectral bleed-through (SBT) parameters, from which final FRET intensities and efficiencies were calculated with the Image J plugin PixFRET ^120^ (version 1.8.0_202). FRET efficiencies were plotted and color-graded with high FRET efficiencies in red and low efficiencies in black [**Fig2F, 2G, suppl. Fig3F**].

### LD Proteolysis

LDs were subjected to trypsin digestion according to a published protocol ^121^. 150 µL of purified LD slurry (∼ 1 µg/µL of total protein; measured with a BCA assay kit (Pierce #23227)) was incubated with 0.016 % w/v of trypsin (Sartorius #03-054-1B) at 37 °C for 90 min. Reactions were gently mixed every 10 min. Following, trypsin was inhibited with 100 μg/mL trypsin inhibitor (Sigma #T9128) for 30 min at 37 °C. Digested LDs were collected by centrifugation at 16000 × g for 3 min and resuspended to a volume of 150 µL in PBS, which was then divided to aliquots of 20 µL. Equal amounts of purified, untreated LD slurries were used as control. Digested and undigested LD aliquots were then mixed with SDS sample buffer, separated on commercial 4-18 % SDS gradient gels by electrophoresis and analyzed by WB with antibodies against Plin2, cPLA2, ATGL and Rab5 (see **Table 2** for details) ^45, 122, 123^. Alternatively, digested and undigested LD aliquots were mixed with fluorophore-labeled wt αSyn to a final protein concentration of 1 μM and the mixture was incubated at 37 °C for 2 h on a shaking platform. LDs were transferred to imaging chambers and protein-LD interactions were detected by fluorescence imaging on a on a Nikon Ti2 CSU-W1 with a 100x objective as described above. [**suppl. Fig3A**]

### Lipid and Cholesterol Inhibitors

Details of inhibitors used in this study are outlined in **Table 3**. Specifically, biosynthesis of lysophospholipids (LPLs) was blocked with inhibitors against calcium-dependent and calcium-independent cytosolic phospholipase A2 (cPLA2), which catalyzes the hydrolysis of phospholipids to arachidonic acid and LPLs, i.e., methyl arachidonyl fluorophosphonate (MAFP, at 10 μM) ^124^, bromoenol lactone (BEL, at 10 μM) ^125^, and pyrrolidine-2 (Py-2, at 1 μM) ^126^. Triacylglyceride (TAG) biosynthesis was inhibited with a broadly-acting antagonist of long-chain acyl-CoA synthetases (ACSLs), which mediate ATP-dependent ligation of fatty acids (FAs) to acetyl-CoA, a key step in the production of DAGs and TAGs, i.e., Triacsin C (TriC, at 5 μM) ^127^, or inhibitors against acyl-CoA:diacylglycerol acyltransferase 1 (DGAT1), i.e., T863 at 20 μM ^128^, and acyl-CoA:diacylglycerol acyltransferase 2 (DGAT2), i.e., PF-06424439 at 10 μM ^129^, which catalyze the final step of DAG to TAG conversion. LD cholesteryl-ester (CE) contents were altered with antagonists of lysosomal cholesterol transport, either by using the general lysosome inhibitor Bafilomycin A1 (BafA1, at 100 nM)^130^ or the specific cholesterol-transport inhibitor U186666A at 20 μM ^131^. For BafA1 rescue experiments, 100 μM M*β*CD-conjugated cholesterol was added to BafA1-inhibited cells (100 nM) for 16 h at 37 °C. CE biosynthesis was blocked with Avasimibe (at 20 μM), which directly inhibits cholesterol esterification by Acyl-CoA:cholesterol acyltransferase (ACAT) ^132^. For all inhibitor experiments, A2780 cells electroporated with 200 μM αSyn were grown on 18 mm fibronectin-coated coverslips at a density of 0.5 × 10^6^ cells for 5 h under standard incubator settings (37 °C, 5 % CO_2_) for recovery. Following, inhibitors were applied in fresh media for another 16 h before cells were fixed and processed for IF analysis. [**Fig3A, suppl. Fig4, suppl. Fig5**]

**Table 3:**
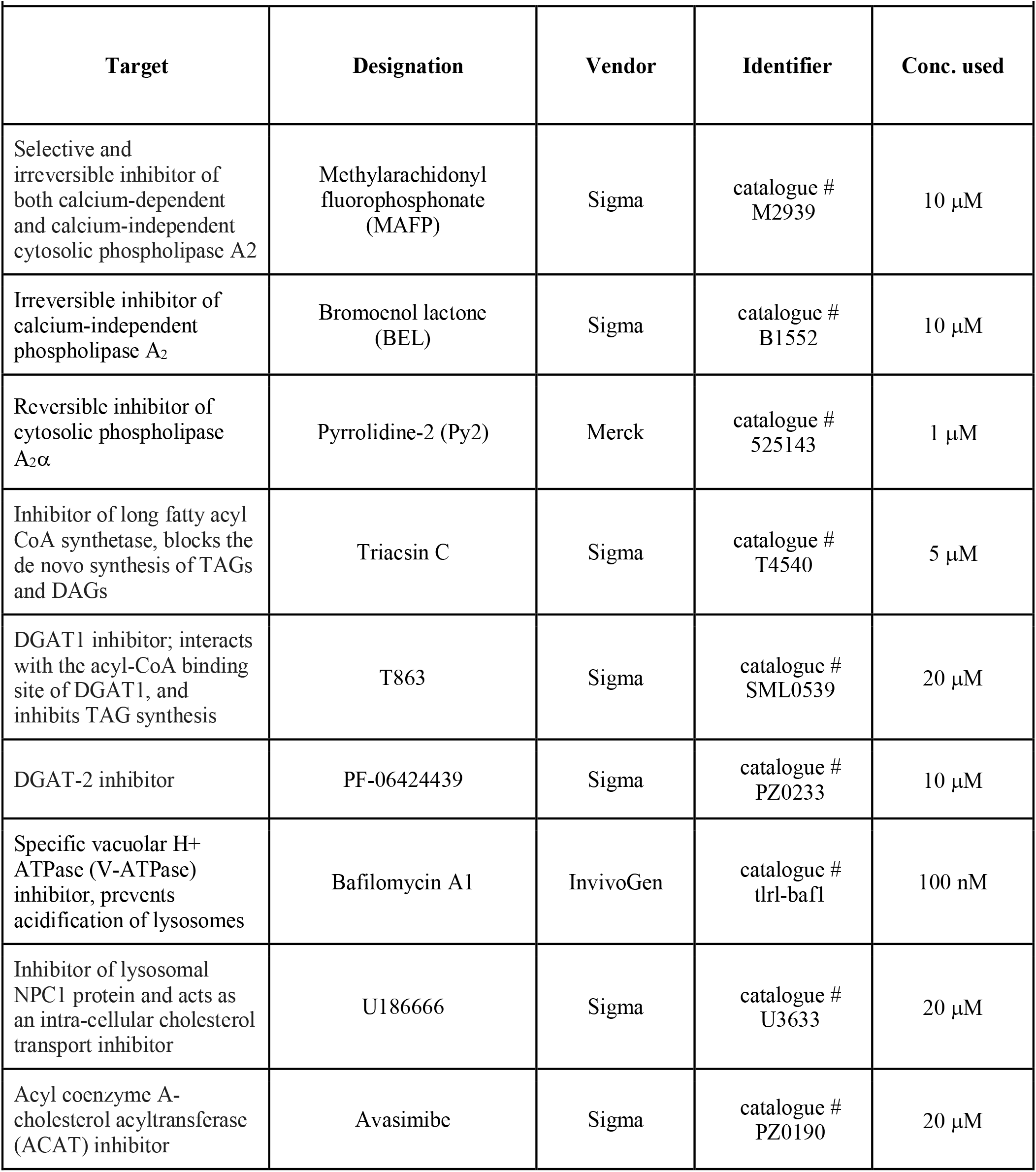
Inhibitors.

### Cholesterol (CHOL), Cholesteryl-Ester (CE) and Triacylglyceride (TAG) Quantification

LD CHOL/CE quantification was carried out with a commercial, colorimetric assay kit (Abcam #ab65359) according to the manufacturer’s instructions and as published ^133^. LDs were purified from A2780 cells grown in CM, or supplemented with OA, PA, CHOL as outlined above, or under starvation conditions in serum-free RPMI without glucose. Purified LD slurries (∼1 µg/µL of total protein; measured with a BCA assay kit) were resuspended in chloroform:isopropanol:NP-40 (7:11:0.1) and vortexed to extract lipids. The suspension was centrifuged at 15,000 × g for 5 min at RT to separate the lipid-containing organic phase. Lipid fraction was initially air-dried at 50 °C to remove chloroform and later vacuum dried for 30 min to remove traces of organic solvent. Resulting pellets were resuspended in 200 μL cholesterol assay buffer. To determine the concentration of *free* CHOL, 2 μL of colorimetric cholesterol dye and enzyme were added to 50 μL of this solution and the volume was adjusted to 100 μL with assay buffer. The mixture was incubated at 37 °C for 60 min in the dark, before absorbance at OD_570_ was measured on a microplate reader (Bio Tek Cytation 5, Agilent). Defined cholesterol standards were processed identically to generate a reference curve. To determine the concentration of *total* CHOL (including CE), 2 μL of colorimetric cholesterol dye and 2 μL of cholesterol esterase enzyme (converting CE to CHOL) were added to 50 μL of the original solution. Further processing was identical to the routine to determine free CHOL. CE content was determined as [CHOL*_total_*] – [CHOL*_free_*]. In practice, [CHOL*_free_*] of analyzed LDs was negligible and [CHOL*_total_*] corresponded to [CE]. Individual measurements were carried out as 2 technical replicates for each batch of purified LDs. To compare individual batches of LDs, we referenced determined amounts of CE (in ng) to the total protein concentration in the original 50 μL slurry. Final values were determined in ng of CE per μg of protein. LD TAG contents were determined using a commercial triglyceride assay kit (Abcam #ab65336) according to the manufacturer’s instructions and as published^134^. Briefly, 150 µL of purified LD slurry (∼1 µg/µL of total protein; measured with a BCA assay kit) was adjusted to 5 % NP 40 and incubated in a water bath at 100 °C for 5 min before cooling to RT. The boil-cool cycle was repeated thrice, and the suspension was centrifuged at 16,000 × g for 5 min at RT. 100 µL of supernatant containing extracted TAG was transferred to a fresh tube and diluted 10-fold with ultra-pure water. For TAG quantification, 50 μL aliquots were removed, colorimetric triglyceride dye/enzyme (2 μL) was added and the volume was adjusted 100 μL with assay buffer. Mixtures were incubated at RT for 60 min in the dark and absorbance at OD_570_ was measured on a microplate reader (Bio Tek Cytation 5, Agilent). For each LD sample, TAG contents were calculated with respect to triglyceride standard curves generated using reference solutions and normalized to total protein concentration measured by BCA assays. CE and TAG ratios were independently confirmed on 2 batches of individually prepared LDs as biological replicates. [**Fig3B, 3C**]

### Yeast LD Purification, Analysis and Imaging

LDs from *Saccharomyces cerevisiae* strains *are1*Δ*are2*Δ (devoid of sterol esters) and *dga1*Δ*lro1*Δ (devoid of triacylglycerols) were isolated according to ^135^. Yeast cells were grown in rich YPD medium containing 2 % glucose (Roth, #HN06.1), 1 % yeast extract (Sigma #13885), 2 % peptone (Sigma #83059) to late exponential/early stationary phase at 30 °C, harvested and spheroplasts were prepared by zymolyase (T20; Amsbio #120491-1) treatment ^136^. Purified spheroplasts were disrupted with a dounce homogenizer in 10 mM MES/Tris buffer (pH 6.9), 0.2 mM EDTA containing 12 % Ficoll (Sigma #F4375). Lysates were separated by successive gradient ultracentrifugation and LDs were obtained at high purity ^137^. Thin layer chromatography (TLC) was performed to visualize yeast lipids. Yeast strains *are1*Δ*are2*Δ, *dga1*Δ*lro1*Δ and wild type (BY4741) were grown to late exponential/early stationary phase, harvested, washed once with water and frozen at -80 °C for 1 hour. Cells were disrupted by glass bead-beating (4 times 30 Hz, 30 s at 4 °C) in TNE buffer (50 mM Tris/Cl, 150 mM NaCl, 5 mM EDTA, pH 7.4) containing 1 mM PMSF (Sigma #P7626). Cell debris and unbroken cells were removed by centrifugation at 500 × g, for 5 min at 4 °C. Proteins in supernatant cell lysates were precipitated with trichloroacetic acid (TCA) at a final concentration of 10 %, solubilized in 0.1 % SDS, 0.1 M NaOH at 37 °C. Total protein concentration was measured as described in ^138^ with bovine serum albumin (BSA) as standard. Lipids from lysates corresponding to 1 mg of total protein were extracted according to ^139^ and separated by TLC with petrol ether:diethyl ether:acetic acid (70:30:2) as solvent. To visualize lipids, TLC plates were developed in 0.63 g MnCl_2_ -4H_2_O, 60 mL H_2_O, 60 mL ethanol, 4 mL concentrated sulfuric acid, dried and heated to 105 °C for 30 min. Cholesteryl-ester and triacylglycerol were used as input controls [**Fig3E**]. To image αSyn-LD interactions, 100 μL of freshly prepared yeast LDs were incubated with AL488-labeled αSyn at a protein concentration of 1.2 μM at 37 °C on a shaking platform. From each of the LD mixtures, 1 µL aliquots were mounted on mPEG-coated, prewarmed coverslips (37 °C). Images were acquired on a Leica SP5 confocal microscope (Leica Microsystems GmbH, Germany) with spectral detection and a 63x NA 1.4 oil immersion objective. AL488 fluorescence was excited at 488 nm, emission was detected between 500-550 nm. Fluorescence and transmission images were acquired simultaneously. Imaging was performed within 5 min of sample loading onto coverslips for each sample. [**Fig3E**]

### Artificial LDs (aLDs)

For preparing artificial LDs (aLDs) of cholesterol oleate (CO, Sigma # C9253), CO was liquified by heating to 50 °C in a water bath. 5 µL of liquified CO was vortexed for 10 s and sonicated for 10 s in 70 µL of HKM buffer at 50 °C (50 mM HEPES, 120 mM potassium acetate, 1 mM MgCl_2_, pH 7.4 and 275 ± 15 mOsm) and then cooled to RT. Triolein (TG, Sigma # T7140 and NBD-triolein (NBD-TG, Setareh biotech #6285) aLDs were prepared by mixing TG and NBD-TG in a 500:1 molar ratio. 5 µL of TG/NBD-TG were vortexed for 10 s followed by sonication for 10 s in 70 µL of HKM buffer at RT. To prepare phospholipid (PL) coated aLDs, a mixture of DOPC:DOPE (Avanti Polar Lipids, #850375 and #850725) (70:30 mol%) in chloroform was dried under a gentle flow of argon gas. Liquified CO or NBD-TG was then added to dried PL to a molar ratio of 0.1%. Mixtures were sonicated for 10 min at 50 °C and the subsequent emulsification process was carried out as described for preparing PL-free aLDs. For interaction assays, TG and CO aLDs with or without PL coats were prepared as described above and mixed. AL594 labeled αSyn was added to this mixture to a final protein concentration of 1 μM. Protein-aLD mixtures were incubated at 37 °C on a shaking platform for 1 h before samples were imaged by confocal fluorescence microscopy [**Fig3F**]. For αSyn and Caveolin1 (Cav1) competition assays, CO aLDs were prepared as described above and incubated with commercially obtained NBD-tagged-Cav1 (NBD-Cav1, amphipathic helix Cav1 residues 159–178, i.e., NBD-LFEAVGKIFSNVRINLQKEI, Peptide 2.0, USA) at 1 μM at 37 °C for 30 min. NBD-Cav1 binding to CO aLDs was confirmed by fluorescence microscopy. Following, 1 μM AL594-labeled αSyn was added to the bulk solution at 37 °C. Displacement of NBD-Cav1 from aLDs by AL594-labeled αSyn was monitored by time-resolved fluorescence imaging on a confocal microscope. [**Fig3G**]

### siRNA Knockdown

Knockdown of *a*Syn in A2780 cells was performed as described ^19^. Commercial siRNA mixtures against human *a*Syn (Dharmacon, ON-TARGET plus human SNCA, #L-002000-00-0005) and a non-targeted control (#D-001810-10-05) were used. A2780 cells were seeded at a density of 6 × 10^5^ cells and transfected with 1.7 µg of the respective siRNA mixtures using Lipofectamine 3000 according to the manufacturer’s instructions. Following transfection, cells were grown for 48 h in CM before analysis. Control and αSyn knockdown cells were subjected to starvation as outlined above. Following siRNA treatment or starvation, A2780 cells were imaged by phase-contrast microscopy on a Nikon Eclipse TS 100 microscope and cells were fixed for IF analysis. Alternatively, cells were harvested by trypsinization after siRNA transfection and/or starvation and Trypan Blue (Sigma #T8154) staining was used to determine cell viability on an automated cell counter (BioRad). Cells were then pelleted by centrifugation and processed for WB. Bar graphs show mean ± SD from three independent experiments (biological replicates). [**Fig4D, 4E, suppl. Fig8A, 8B**]

### LD Counting

To determine LD numbers, cells were imaged on a Nikon Ti2 CSU-W1 with a 100x objective and analyzed using Fiji ^118^. Cell outlines were identified and drawn manually based on phase contrast or αSyn IF images. Individual cells were defined as ROI (region of interest). LDs in each ROI were first identified using BODIPY signals and processed as follows: 16-bit images were converted to 8-bit and thresholds were set using an auto-threshold function as outlined in ^140^. Following, images were converted to binary masks and adjacent objects were separated using the watershed function. The Fiji particle counting routine was applied to determine LD numbers and dimensions in each ROI. The same data extraction was repeated for every image in the data set. Blots correspond to 110–120 combined ROIs (cells) from three independent experiments (biological replicates). [**Fig4E, Fig5A, 5B, 5D, suppl. Fig8E**]

### Cell Lysates

Lysates of A2780 and HEK cells were prepared by detaching ∼5-10 × 10^6^ cells with trypsin/EDTA (0.05% / 0.02%), collected by centrifugation at ∼200 × g for 5 min at 25 °C. Sedimented cells were washed once with PBS, counted on an automated cell counter (BioRad) and pelleted again by centrifugation. After resuspending cells in PBS with cOmplete proteinase inhibitor cocktail (Roche #04693132001), slurries were adjusted to 2 × 10^7^ cells/mL and lysed by three freeze-thaw cycles. Lysates were cleared by centrifugation at 16000 × g for 30 min. Supernatants were collected and total protein concentration was measured with a BCA assay kit (Pierce #23227). 50 µg of protein per lane [**suppl. Fig8C**] or 30 µg of protein per lane [**suppl. Fig9A, 9B**] were applied and separated by SDS-PAGE and transferred onto PVDF membranes for WB.

### Western Blotting (WB)

Cell lysates and recombinant protein samples were boiled in Laemmli buffer for 10 min before SDS-PAGE separation on commercial, precast 4-20 % gradient gels (BioRad #4561095). Recombinant N-terminally acetylated αSyn at indicated concentrations was loaded as reference input. Proteins were transferred onto PVDF membranes (Biorad #1704157) and fixed with 4 % (w/v) PFA in PBS for 1 h according to ^141^. Membranes were washed twice with PBS, twice with TBS, 0.1 % tween 20 (TBST) and blocked in 5 % dry milk-TBST for 1 h. After blocking, blots were incubated with primary antibodies overnight at 4 °C. Membranes were washed and incubated with horse-radish peroxidase (HRP)-conjugated secondary antibodies for 1 h (at RT). Membranes were developed using the SuperSignal West Pico Plus reagent (ThermoScientific #34579) and luminescence signals were detected on a BioRad Molecular Imager. Intensities of *a*Syn and *b*-Actin bands were quantified using the ImageLab software (BioRad). *a*Syn reference input was used to generate standard curves. αSyn concentrations in each lysate sample were calculated with respect to αSyn standard curves and normalized to *β*-Actin signals. Bar graphs show mean ± SD from two independent experiments (biological replicates). For cell lysate samples, *a*Syn intensities were normalized according to *b*-Actin signals and *a*Syn lysate concentrations were calculated with respect to standard curves. Standard deviations were calculated from 2 biological replicates. [**suppl. Fig8A, 8C, suppl. Fig9A, 9B**]

### Statistical Analysis

Tukey box plots show median values (center lines) with box dimensions representing the 25^th^ and 75^th^ percentiles. Whiskers extend to 1.5 times the interquartile range from the 25^th^ and 75^th^ percentiles. For box plots, data points considered ‘outliers’ were determined based on the criteria defined in the outlier test by the ROUT method ^142^ and omitted. Analysis of variance (one-way ANOVA) tests with Bonferroni’s post-tests ^143, 144^ were used to determine the statistical significance of experiments with more than two samples, whereas Student’s t tests were performed to assess statistical differences between samples. Significance is given as ns > 0.05, ∗ p < 0.05, ∗∗ p < 0.01, ∗∗∗ p < 0.001. Graphs were generated and statistical analyses were performed in GraphPad Prism (version 9.4.1).

## Acknowledgments

We thank Martin Lehmann (Cellular imaging, FMP-Berlin) and Yoseph Addadi (Life Sciences Core Facilities, Weizmann Institute of Science) for excellent maintenance of imaging facilities and valuable discussions along the project. We are grateful to Dr. Shira Albeck and the Protein Purification Unit (Life Sciences Core Facilities, Weizmann Institute of Science) for the preparation of individual batches of recombinant aSyn and to Dr. Kota Miura (The Network of European Bioimage Analysts, NEUBIAS) for his insight in interpreting FRAP data and discussing FRAP/FRET models. FRET and FRAP experiments were acquired at the Advanced Optical Imaging Unit, de Picciotto-Lesser Cell Observatory Unit at the Moross Integrated Cancer Center Life Science Core Facilities at the Weizmann Institute of Science. We additionally thank Profs. Hagen Hofmann and Atan Gross (Weizmann Institute of Science), Andres Binolfi (CONICET, Rosario Argentina), Stefano Vanni (University of Fribourg), Paola Picotti (ETH Zurich), Julia Mahamid (EMBL Heidelberg), Melissa Birol (BIMSBI Berlin) and Cedric Eichmann for highly valuable inputs and expertise throughout the project. We are grateful to Drs. Veijo Tuomas, Verneri Salo (EMBL Heidelberg) and all members of the group for carefully reading the manuscript. PS acknowledges funding by the European Research Council (ERC) Consolidator Grant NeuroInCellNMR (647474). Work in the Selenko laboratory is supported by the Willner Family Foundation.

## Notes

### Competing Interest Statement

The authors have declared no competing interest.

